# Subjectivity of time perception alters choice preference for future rewards through fronto-striatal value signal dynamics

**DOI:** 10.1101/2022.03.16.484555

**Authors:** Reiko Shintaki, Daiki Tanaka, Moe Okayasu, Shinsuke Suzuki, Takaaki Yoshimoto, Norihiro Sadato, Junichi Chikazoe, Koji Jimura

**Author notes:** Correspondence should be addressed to: Koji Jimura, Ph.D., Department of Biosciences and Informatics, Keio University, 3-14-1 Hiyoshi Kohoku-ku, Yokohama, 223-0061, Japan. The authors declare no conflict of interest.

## Abstract

Our preference for a reward depends on the time of delay for its delivery. Here we show that changes in the external environment can manipulate the perceived duration of time, which alters the formation of the choice preference through value signals in the cortical and subcortical brain regions. Humans anticipated a real liquid reward delayed by tens of seconds, during which colors of visually presented panels gradually changed. The color-change delay was perceived as shorter than a control color-constant delay. Interestingly, participants with greater perceptual bias of the delay showed stronger preference for rewards with the color-change delay. The ventrolateral prefrontal cortex (vlPFC) and ventral striatum (VS) showed dynamic neural signatures of value components that were modulated by subjectively perceived duration. Crucially, these effects were specifically observed while a future reward was anticipated, and moreover, the vlPFC activity was weaker in participants with greater bias in the duration perception. These results demonstrate that subjective time experience leads to biased choice preference of delayed rewards, which is regulated by dynamic value signals of future rewards and accurate perception of the external world in the vlPFC-VS systems.

---

In our daily life, we often perceive different durations of time as longer or shorter, even if their objective durations are identical. For example, on a round trip journey, the duration of the return trip is perceived as shorter than that of the outbound trip, even though their routes and durations are objectively identical^1–3^. This flexible, elastic subjectivity in the perception of time has been observed in various behavioral and cognitive situations, with perceived durations varying depending on the temporal frequency of change in external events^4–6^, environmental complexity^7, 8^, and expectation of duration^1, 9^.

Everyday decision-making often includes choices between options that differ in terms of on multiple factors. One critical factor is time to outcome. In reward-related decision-making, a reward delivered in a remote future is devalued compared to another that is available sooner, if the rewards are otherwise identical^10, 11^. Such valuation has been examined in intertemporal decision-making involving evaluation of future outcomes that vary both in their magnitude and time of delivery^12, 13^. In standard experimental paradigms, behavioral agents make a choice between a larger delayed reward or a smaller immediately available reward^14–18^. The value of the delayed reward is then estimated as an equivalence of the immediate reward^13, 19–22^. The subjective value is known to decrease with longer delay time, a phenomenon referred to as delay discounting^23–26^.

Given the subjectivity of time perception^4–9^, valuation of the delayed reward should also depend on the subjectively perceived duration of the delay. Time discounting of a future reward would then be mediated accordingly. More specifically, a reward option with a delay that is perceived as shorter would be preferred to another even if their objective delay durations, reward types, and reward amounts are identical. As such, the subjectivity of time experience may play an important role in intertemporal decision-making.

Decision neuroscience has demonstrated that neural signals represent temporally evolving value components regarding a future reward while the reward is anticipated^22, 27, 28^. In particular, the ventral striatum (VS) shows temporally modulated signals coding the value of a future reward that is discounted as a function of the duration until the reward is delivered; signals in the prefrontal cortex (PFC) reflects the current utility of anticipating future rewards^22, 27–29^. On the other hand, neurophysiological studies have identified sets of neurons that successively fire along with temporally structured experiences^30–36^, suggesting that the experience of time is encoded in coordinated neuronal activity. Thus, a subjectively perceived duration may regulate the dynamic value signals during anticipation of a future reward.

The current study aimed to examine whether the subjectivity of perceived duration modulates the value of delayed rewards. To this end, we designed a novel behavioral experiment that was conducted during functional MRI (fMRI) scanning (Figure 1). Human participants performed a variant of the intertemporal choice paradigm where they received a real liquid reward delayed by dozens of seconds^20–22, 27^ (delayed reward choice task; Figure 1 and Supplementary Figure 1). A key experimental manipulation was that to bias perceived delay duration, participants were presented with a set of colored panels (Figure 1a *bottom*) whose colors continuously and gradually changed (color-change delay), or remained unchanged (color-constant delay) through the delay while the participants awaited a reward (Figure 1b). They first received two delayed rewards with the color-change and -constant delays, with the goal of enabling them to form a choice preference for the rewards through direct experience^27^ (formation trials). Then, they chose which of the two alternatives they preferred (choice trial; Figure 1c *bottom left*). To estimate the subjective duration of the color-change delay, they performed another task in which they were presented with the same color-change and -constant delays without receiving a liquid reward, but they instead judged which of the delays seemed longer (duration judgment task; Figure 1c *bottom right*). To examine whether subjective duration was reflected in brain activity while awaiting a future reward, we examined the temporal dynamics of brain activity reflecting value-related signals during the delay periods^13, 22, 27–29, 37–40^, and adjusted the signals by the subjective duration of the delay. Then, we explored brain regions showing dynamic brain activity modeled by the value components.

**Figure 1.**
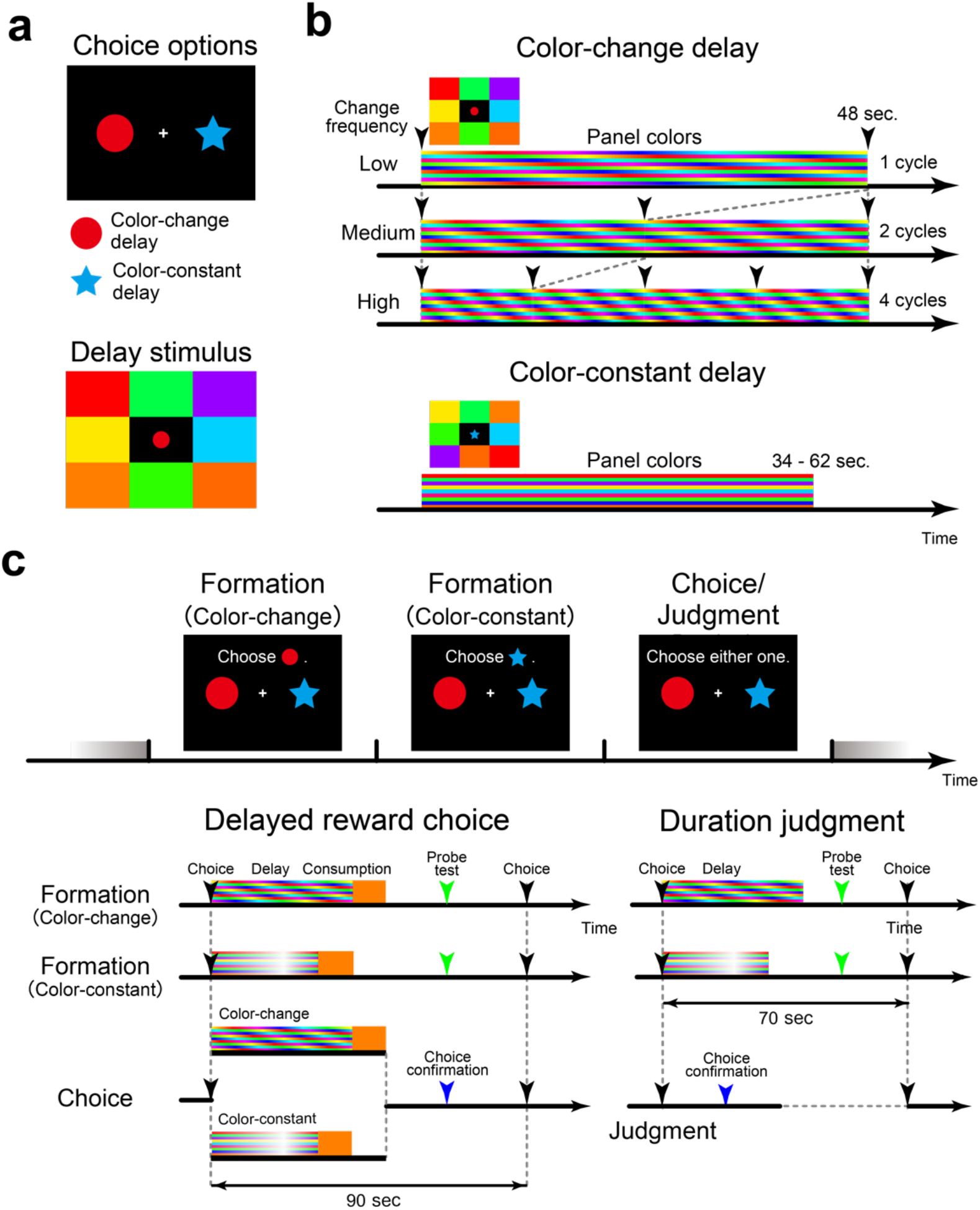
Study design. **a**, *Top*, choice situation. Pictures of two items were presented on the screen, and both pictures corresponded to a real liquid reward delayed by tens of seconds. The colors of eight colored panels that were presented peripherally either changed (red circle) or did not change (blue star) during the delay period. Each picture was used in only one trial block. *Bottom*, delay-period stimulus. While participants waited for a reward, the eight color panels were presented and the picture choice was presented centrally. **b**, During the color-change delay, the colors were changed gradually and continuously (*top*). The color change frequency was low, medium, or high. The panel colors were constant during the color-constant delay period (*bottom*). The horizontal bars represent the colors of the panels. **c**, Participants first experienced two delayed rewards, one with the color-change delay and the other with color-constant delay (*top*). In the reward choice task (*bottom left*), they received and consumed a real liquid reward after the delay period (formation trial). Then, they made a choice between the two picture choices based on their past experiences in the formation trials (choice trial). These three trials (a block) were performed in triplicate. In the duration judgment task (*bottom right*), after they experienced two delay periods, they judged which of the delays was longer.

## Results

We first analyzed the subjective duration of the color-change delay in the duration judgment task. The frequency of the change was low, medium, or high (Figure 1b *top*). For each frequency, the duration of the color-constant delay was adjusted based on prior judgments to estimate the subjective duration of the color-change delay (Supplementary Figure 1b; see Methods). Then, we defined the subjective duration of the color-change delay as equivalent to that of the color-constant delay^20–22, 27^. We observed that subjective duration was shorter in the color-change delays than in the color-constant delay [t(26) = 2.2; P < .05; Figure 2a], and high-frequency color-change delays were perceived as shorter than low-frequency ones [linear trend: t(26) = 3.3; P < .01].

**Figure 2.**
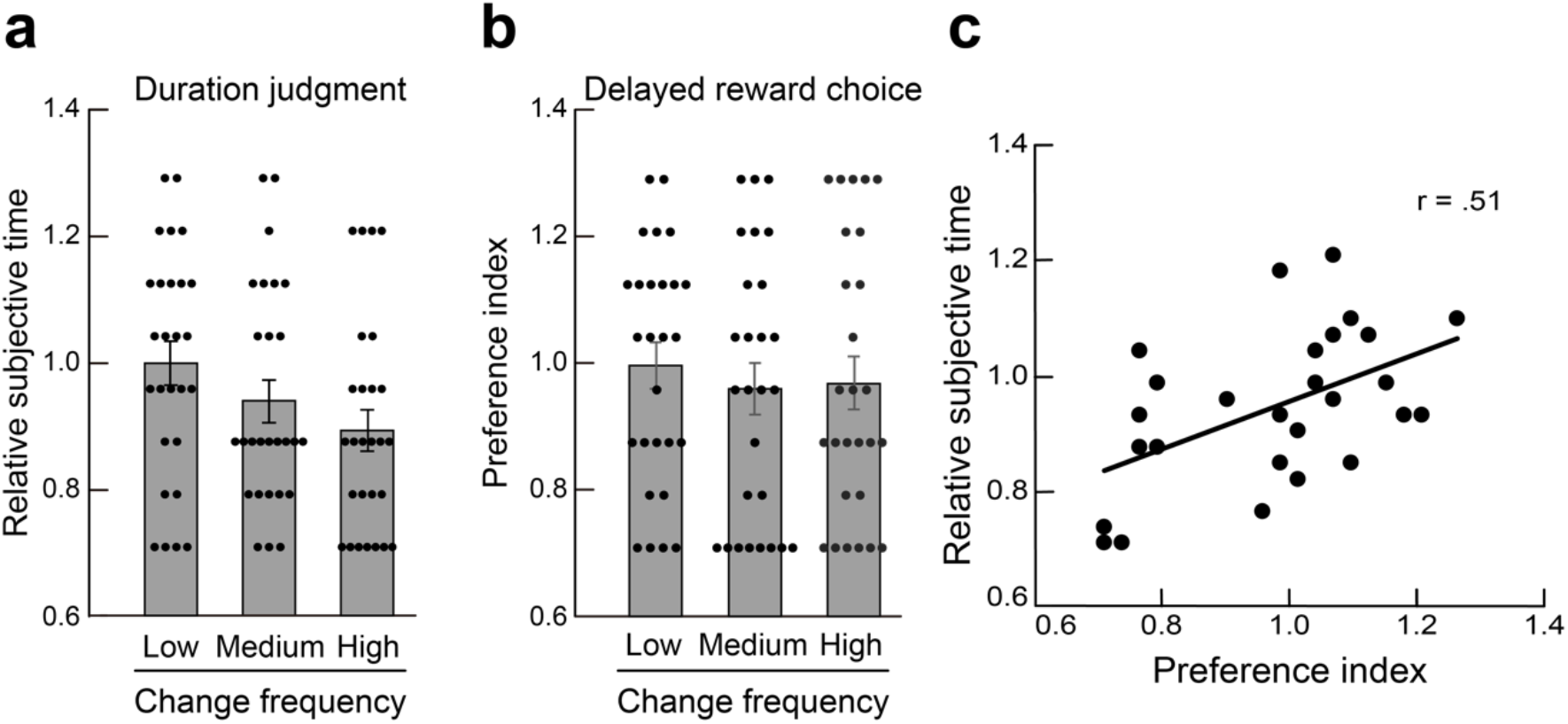
Behavioral results. **a**, Subjective duration of the color-change delay as an equivalence of a color-constant delay estimated in the duration judgment task. The subjective duration was normalized by the duration of the standard delay duration (48 s). Thus, a value less than 1 indicates that the color-change delay was perceived as shorter than a 48-s color color-constant delay. Error bars indicate standard errors of the mean across participants. The dots represent individual data. **b**. Choice preference index of the color-change delay reward estimated in the reward choice task. Note that a smaller value of the preference index indicates a strong preference (greater valuation) for the color-change delay reward. **c**, A scatter diagram of the relative subjective duration and preference index. Each plot denotes one participant. **: P < .01.

In the reward choice task, a real liquid drink was used as a reward, and was delivered after the delay period^20–22, 27^. The drink and its amount were identical for the rewards in the color-change and -constant delays, and the only differences were the objective delay duration and whether the panel colors changed. We quantified the choice preference for the delayed reward with the color-change delay. The duration of the color-constant delay was adjusted based on prior choice, as in the duration judgment task (Supplementary Figure 1a; see Methods). Then, the preference index was defined as an equivalent duration of the color-constant delay. A preference index less than 1 indicates that the choice preference for a color-change reward is equal to that to a color-constant reward with shorter objective delay duration. Thus, a smaller preference index can be interpreted as a stronger preference to a color-change reward because the subjective duration of the color-change delay is perceived as being shorter (Figure 2a). Participants did not show a significant preference for the color-change reward [t(26) = 0.8; P = .42; Figure 2b], although the choice preference was biased toward the color-change delay in all color-change frequencies.

Nonetheless, if the subjective perceived duration modulates the formation of the choice preference, the modulation of choice preference should consistently appear across individual participants. To test this possibility, we examined the correlation between the preference index and subjective duration. For each participant, preference indices and subjective durations were averaged across color-change frequencies, and Pearson’s correlation coefficient was calculated across participants. We observed a strong positive correlation between them [r = .51; t(25) = 3.0; P < .01; Figure 2c]. This positive correlation indicates that individuals who perceived the color-change delays as shorter showed stronger choice bias toward the reward with the color-change delay. We note that it is reasonably conceivable that overall preference for the reward with the color-change delay was weak (Figure 2b) because the choice preference for delayed rewards does not depend solely on delay duration^13, 20, 29^ (see also Discussion). Thus, our results highlight that the subjective duration of the delay is one of the crucial factors influencing the choice preference of delayed rewards.

### Dynamic value representations during anticipation of a future reward

We so far demonstrated that the subjectively perceived duration of the color-change delay was shorter than that of the color-constant delay, and that this was associated with the formation of choice preference for the delayed rewards. The question then arises as to whether the subjective experience of the delay period is coded in brain activity while a future reward is anticipated. To address this issue, we the examined temporal characteristics of brain activity during the delay period of the formation trials. In particular, we focused on the representation of temporally evolving value components regarding a future reward as forecasted and demonstrated in previous studies^13, 22, 27–29, 37–40^. Two temporal components were modeled. One was the anticipatory utility (AU) model, which reflected the utility of waiting for a future reward^40^, and the other was the upcoming future reward (UFR) value model, which explained the dynamic increase in future reward value as its delivery approached^13, 38, 41^. The AU model showed a peak value at the beginning of the delay and a gradual decrease along the course of delay, while the UFR model had inverse temporal characteristics^13, 22, 27, 28, 38–41^ (see Methods).

Importantly, prior to every formation trial, participants were naive to the upcoming delayed reward because they had no experience with the choice options, which were indicated by unique pictures (Figure 1a). However, based on past trials they had experienced, it should have been possible to expect when a future reward would become available. Thus, the current analysis assumed that prior to each formation trial, participants had expectations regarding the delay duration, and the expectation was updated through their experiences in the formation trials^27, 28^.

To implement this idea, the current analysis used the Bayesian inference approach^27, 28, 42^ (see Methods). The expectation of reward outcome was modeled stochastically, based on a probability density distribution of reward outcome occurring in the future. The probability distribution was calculated using a gamma function as a function of the expected delay duration (eq. 1), and the distribution was updated every time participants completed a formation trial (eq. 2).

One important issue in the current experiment is that participants perceived the color-change delay as being shorter than the color-constant delay (see Figure 2a; Behavioral results). Thus, the probability density distribution should be modulated in accordance with the subjectivity of the duration perception. It should be noted that the perceived duration of a delay was estimated as an accumulation of subjective units of time that elapsed from moment to moment^4^. Therefore, the shorter subjective duration of the color-change delay indicates that the subjective course of time elapsed more slowly during the color-change delay than the color-constant delay. Thus, the probability distribution of reward delivery with the color-constant delay should be wider along the temporal axis, because the subjective course of time was slower; accordingly, its peak should be lower.

We introduced this theoretical consideration about the subjectivity of duration perception into our modeling. Specifically, the probability distribution was modulated by the subjective duration estimated based on the duration judgment task, such that the probability distribution of the color-constant delay was multiplicated by the relative subjective duration, and its time constant (i.e., speed of temporal change) was divided by the relative subjective duration (Figure 3a *left* for an example participant; eq. 3; see Methods).

**Figure 3.**
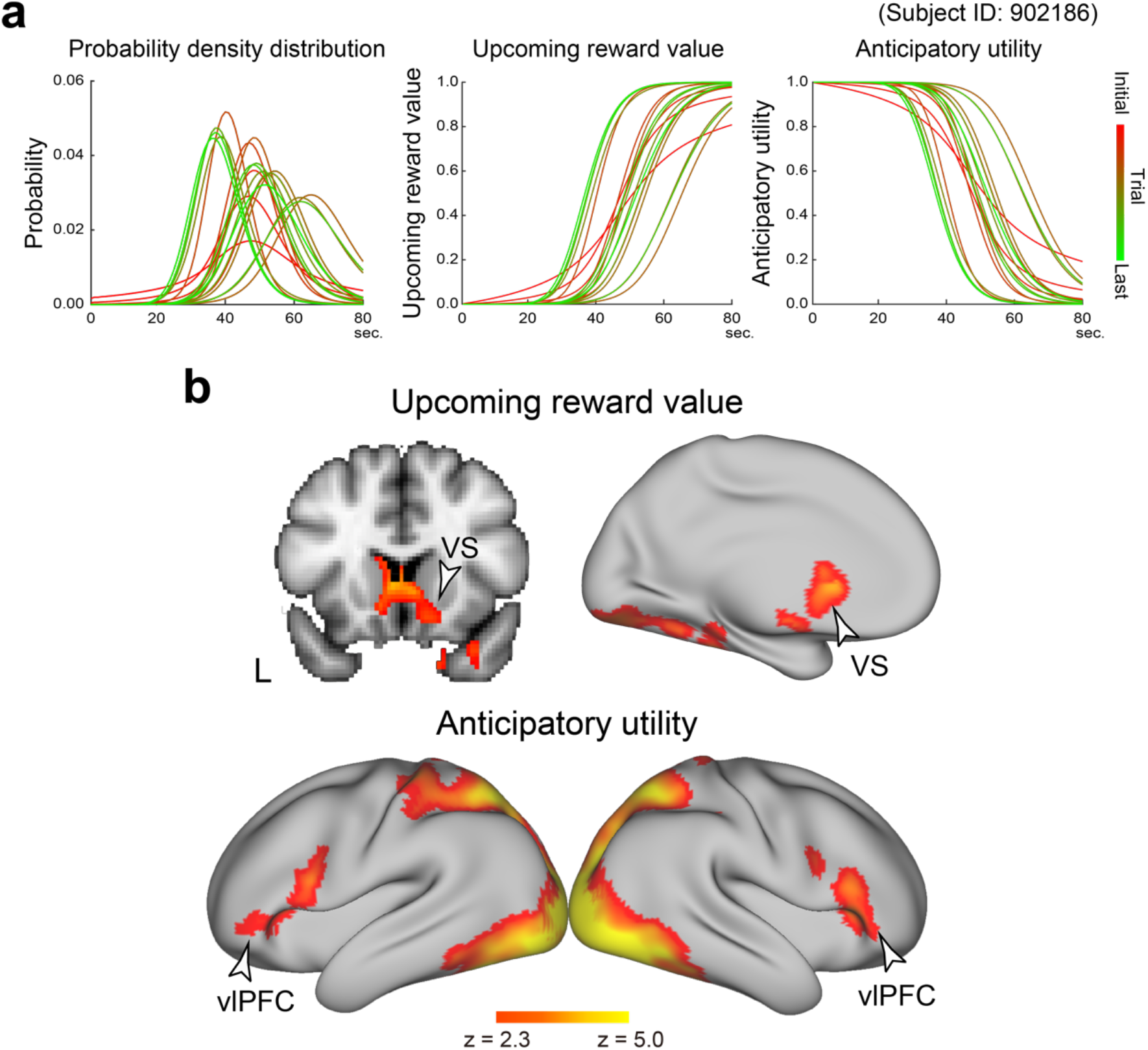
Dynamic value components while anticipating a delayed reward. **a**, *Left*, the probability density distribution for the expected delay was formulated by a gamma function for an example participant (ID: 902186). Distribution parameters were updated based on Bayesian inference according to participants’ experiences in the formation trials. Then, the probability density distribution was adjusted by the subjective duration of the delay. The line colors indicate trial experiences of delayed rewards as shown in the bar on the right. *Middle*, upcoming figure reward (UFR) dynamics were modeled by the accumulated probability from the delay onset until the present time. *Right*, anticipatory utility (AU) was modeled as the inverse of UFR. **b**. Brain regions showing dynamic value signal components during the delay period in the formation trials. UFR effect (t*op*). AU effect (*bottom*). Maps were overlaid on the 3D surface of the standard brain or a coronal section. The color bars indicate significance levels. VS: ventral striatum; vlPFC: ventrolateral prefrontal cortex.

To model UFR dynamics during the delay period, the probability density function was integrated from the start of the delay to the current time (eq. 4; Figure 3a *middle*; see Methods). The idea behind this integration is that, as the delay period was experienced, participants’ expectation of a reward continued to increase, like hazard rates^37, 43^. Thus, the UFR model was formulated as the cumulative probability from the start of the delay to the current time, although participants did not know exactly when the reward would become available. Then, the AU model was defined as 1 – UFR to model inverse dynamics, similarly to prior modeling^22, 27, 28^ (eq. 5; Figure 3a *right*). The UFR and AU models were calculated for each formation trial for each participant, and were convolved with the canonical hemodynamic response function (HRF) (Supplementary Figure 2a). Then, model parameters were estimated based on a standard general linear model (GLM) approach.

Whole-brain exploratory analysis of these temporally dynamic value components identified a region in the ventral striatum (VS) that showed a strong UFR effect (Figure 3b *top*; Supplementary Table 1; color-change and -constant delay collapsed). On the other hand, the AU effect was observed in regions across the brain including the bilateral ventrolateral prefrontal cortex (vlPFC) and posterior parietal cortex (Figure 3b *bottom*; Supplementary Table 1).

If the UFR and AU reflect value representation regarding a future reward, these dynamics signals should be absent during a delay period in which a future reward is not anticipated. To test this possibility, we modeled the dynamic value components during the delay period in the duration judgment task, where a reward was not anticipated, and performed GLM estimations. Such dynamic effects were absent in the VS and vlPFC (Supplementary Figure 3; Supplementary Table 2).

Then, the two dynamic value components were directly compared between the delayed reward choice task and the duration judgment task by contrasting the AU and UFR effects between the two tasks. In the vlPFC, the AU effect was greater in the reward choice task than in the duration judgment task (Figure 4a; Supplementary Table 3). Importantly, during the delay period in the two tasks, identical visual stimuli were presented, but whereas participants anticipated a future reward in the reward choice task, this expectation was not present in the duration judgment task.

**Figure 4.**
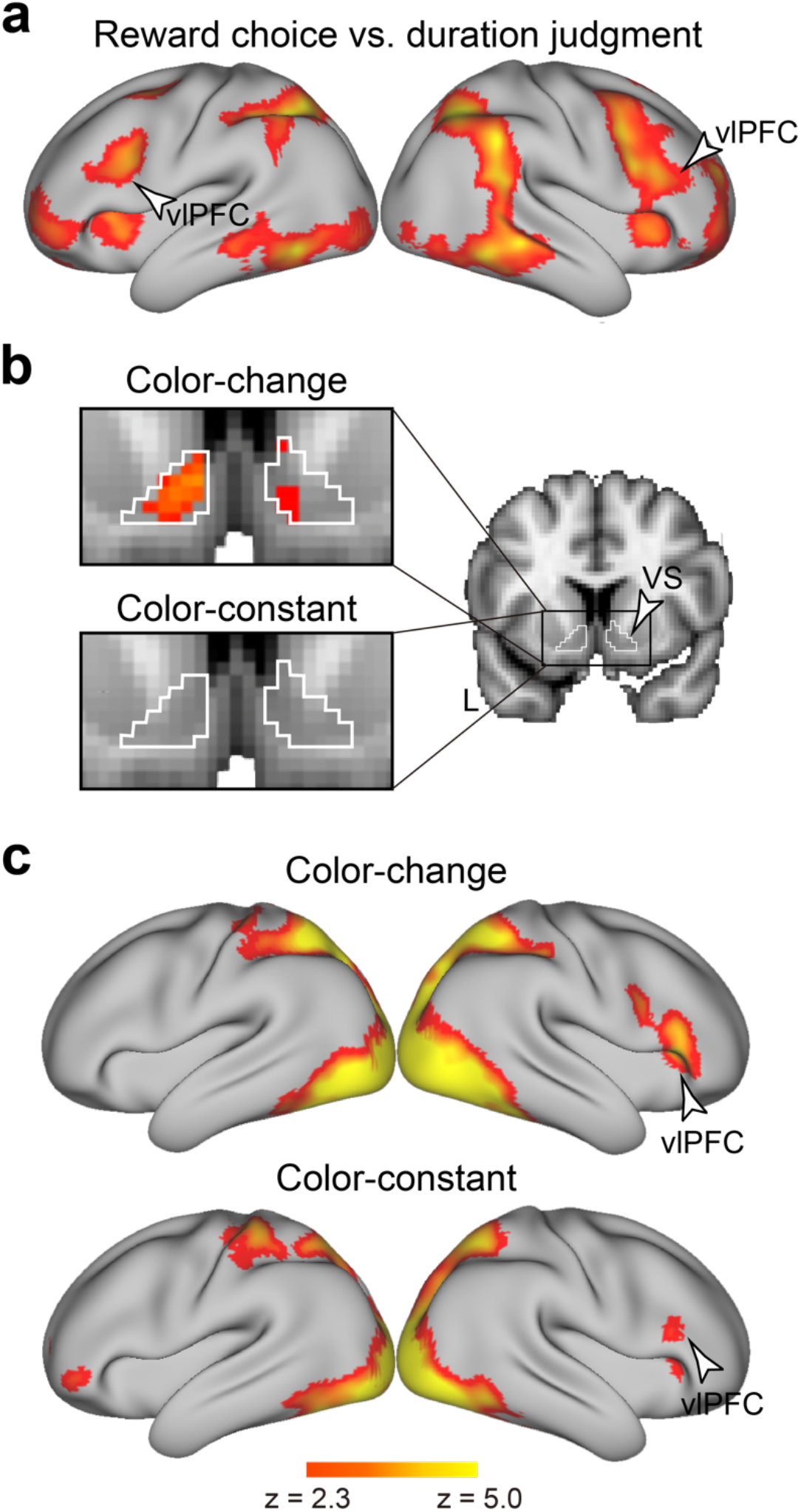
**a**, Brain regions showing greater AU effects in the reward choice task relative to the duration judgment task. **b**, The UFR effects in the VS were estimated separately for the color-change delay (*top left*) and color-constant delay (*bottom left*), and mapped onto 2D-slice anatomical images. The black solid square on the VS (*right*) represents the areas magnified on the left. White lines indicate the anatomically defined VS region. **c**, The AU effects were estimated separately for the color-change delay (*top*) and color-constant delay (*bottom*), and mapped onto the 3D surface of anatomical images. Formats are similar to those in Figure 3b.

Our analysis revealed that the UFR and AU effects occurred specifically during anticipation of a future reward. On the other hand, our behavioral results demonstrated that the subjective duration of the color-change delay was shorter, which was associated with a stronger preference for the reward with the color-change delay. The question then arose whether the UFR and AU effects were consistently observed during the color-change and -constant delays or whether these delays yielded differential effects. To test this possibility, we first estimated the UFR and AU effects for these delays separately. During the color-change delay, the VS and vlPFC showed strong UFR and AU effects, respectively, while these effects were attenuated during the color-constant delay (Figures 4b/c).

Nonetheless, when the UFR and AU effects were directly contrasted between the color-change and -constant delays, differential effects were absent in the prefrontal and striatal regions. No brain regions showed differential UFR effects, and the AU effect in the color-change delay was greater only in the occipital and parietal regions (Supplementary Figure 4). Additionally, when the UFR and AU effects were compared between the two tasks separately for the color-change and -constant delays, the vlPFC showed greater AU signals during the reward choice task in both delays (Supplementary Figure 5). These collective results suggest that during anticipation of a future reward, the VS and vlPFC showed dynamic value signals independently of the changes in the external environment, and the signals became even greater when the external environment changed.

### Dynamic value representations during the delay period of reward choice trials

We next examined the UFR and AU effects during the delay period of the reward choice trial in the delayed reward task. As in prior models^27, 39^, we used linear functions to model the temporal characteristics of the UFR and AU components (Supplementary Figure 2b; see Methods). Whole-brain exploratory analysis identified brain regions showing a strong UFR effect in the VS (Supplementary Figure 6a *left*; Supplementary Table 4) and in the ventromedial prefrontal cortex (vmPFC; Supplementary Figure 6a *right*; Supplementary Table 4). On the other hand, bilateral polar regions in the anterior prefrontal cortex (aPFC) exhibited the AU effect (Supplementary Figure 6b; Supplementary Table 4). These results were highly consistent with those of prior analyses of the UFR and AU effects^22, 27, 28^.

### vlPFC associated with subjective delay duration

Thus far we observed the following: 1) The AU effect in the vlPFC was specifically associated with anticipation of a future reward (Figures 3b *bottom* and 4a), 2) the AU effect in the vlPFC was more prominent during the color-change delay (Figure 4c), 3) the color-change delay was perceived as shorter (Figure 2b), and 4) greater perceptual bias produced a stronger preference for the reward with the color-change delay (Figure 2c).

Given these results, we asked whether the AU-related vlPFC regions were associated with the subjective duration of the color-change delay. To address this issue, we performed an exploratory analysis to determine which regions within the vlPFC that showed AU effect (Figure 3b *bottom*) also exhibited greater activity during the color-change delay than the color-constant delay. In this analysis, we analyzed imaging data during the duration judgment task, not the delayed reward choice task, because the former is more appropriate for evaluating the effect of subjectively perceived duration. Additionally, we examined a block effect with a constant value through the delay, not the dynamic value component of the AU or UFR, because anticipation of a future reward was not a component of this analysis. The analysis revealed that a vlPFC region showed a greater activity during the color-change delay relative to the color-constant delay [(38, 22, 18); 924 voxels; peak t(25) = 6.6; P < .05 FWE-corrected within vlPFC region showing AU effect; Figure 5a *left*].

**Figure 5.**
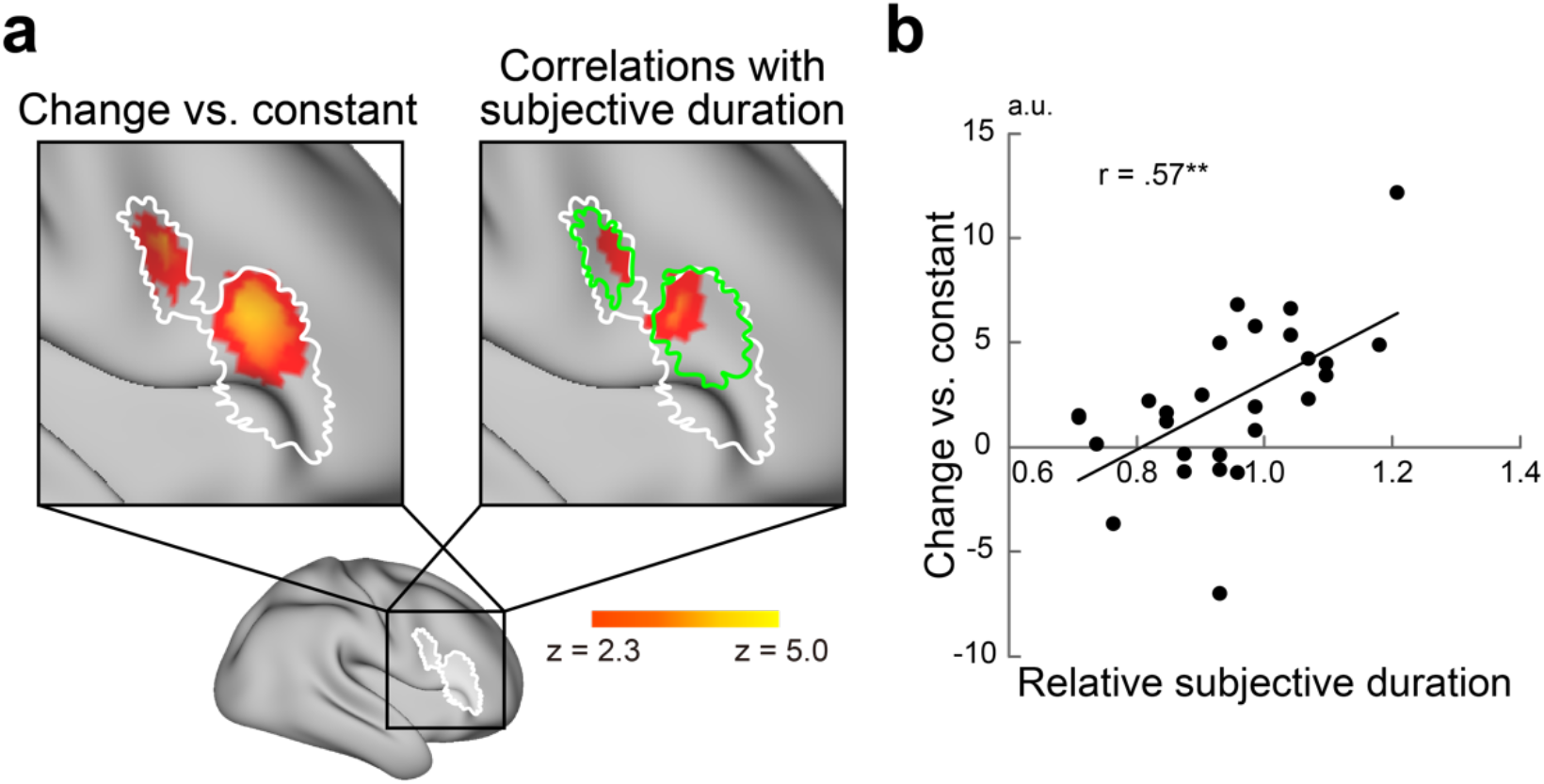
Block-effect activity during the delay period in the duration judgment task. **a**, Brain regions showing greater activity in the color-change delay than in the color-constant delay (*left*). Brain regions showing correlations between relative subjective duration and activity (color-change vs. color-constant) (*right*). The black solid square on a 3D surface map (*bottom*) represents the areas magnified on the top. White lines indicate vlPFC regions showing AU effects during the delay period of the reward choice task. Green lines indicate the activated vlPFC regions in the left panel. Formats are similar to those in Figure 3b. **b**, A scatter diagram of relative subjective duration and brain activity in the vlPFC during the delay period of the duration judgment task. The region of interest in the vlPFC was defined independently of the current data. Each plot denotes one participant. **: P < .01.

To further specify the relationship between vlPFC activity and subjective duration, we performed another exploratory analysis identifying vlPFC regions showing correlations between brain activity (color-change delay vs. -constant delay) and the subjective duration of the color-change delay (Figure 2a). Interestingly, the same vlPFC region showed a strong positive correlation [(48, 22, 16); 151 voxels; peak t(25) = 4.8; P < .05 FWE-corrected within vlPFC region showing AU effect; Figure 5a *right*]. These results demonstrated that this area of the vlPFC showed a conjoint effect of 1) AU during the delayed reward choice task, 2) greater activity during the color-change delay, and 3) correlation between color-change-related activity and subjective duration.

Behavioral analysis showed that the color-change delay was perceived as shorter. Thus, the positivity of the correlation indicated that enhanced shortening (bias) of the subjective duration is associated with weaker vlPFC activity during the color-change delay. Importantly, our result is consistent with a recent study showing that this vlPFC region is associated with accurate time perception^44^. Of note, the reported vlPFC coordinate in the recent study was (48, 24, 14), which was 2.8 mm away from the correlation peak in the current study. We then performed a region of interest (ROI) analysis in which the ROI was defined based on the coordinates reported in the prior study^44^. In this ROI, we observed a strong AU effect during the color-change delay in the reward choice task [t(26) = 3.8, P < .001], greater activity during the color-change delay compared to the color-constant delay in the duration judgment task [t(26) = 3.0, P < .01], and correlation between the activity and subjective time [r = .57; t(25) = 3.5; P < .01; Figure 5b].

## Discussion

The current study manipulated subjective experiences while humans were awaiting a future reward. The subjective duration of the delay time became shorter when they experienced a change of colors, an effect that was enhanced by an increased frequency of the change. Individuals with greater perceptual bias showed a stronger preference for the reward with the color-change delay. The temporal dynamics of the value components during the delay period of the formation trials were modeled such that the dynamic components were modulated by the subjective duration of the delay. The activity in the VS and vlPFC showed dynamic pattern of the UFR and AU models, respectively. Additionally, a greater bias in duration perception was associated with lower vlPFC activity. These results suggest that dynamic vlPFC-VS systems yield choice biases for delayed rewards depending on the subjective experience of a delay while a future reward is anticipated.

### Subjectivity of delay duration and reward value

To estimate the subjective value of a delayed reward, standard intertemporal choice paradigms have adjusted the amount of an immediate or delayed reward^20–22, 45^. In contrast, the current study adjusted the duration of the color-constant delay to estimate the value of a delayed reward with the color-change delay, thus yielding the preference index. This manipulation hypothesizes that the subjective value of a delayed reward is a function of the subjectively perceived duration of delay time.

In the duration judgment task, the subjective duration of the color-change delay was perceived as shorter than that of the color-constant delay. This result allowed us to predict that underestimation of the perceived duration of the color-change delay would increase the subjective reward value with the color-change delay (i.e., a decrease in the preference index). However, this effect failed to reach a statistical significance, although the direction of value modulation was consistent.

We did not consider the attenuated effect unreasonable, because the value of the delayed reward depends on multiple factors, and not solely on delay duration^13, 20, 29^. These factors include reward type^21^, reward amount^13, 20^, and possibly preference of delay period experiences. For example, some participants may perceive the color-change delay as being shorter, but will not prefer changes in stimulus colors *per se*, which may diminish the potential increase of the reward value with the color-change delay. The results of post-task questionnaires supported this possibility.

Nonetheless, the significant positive correlation between the preference index and the subjective duration of the color-change delay (Figure 2c) demonstrates that this duration affects the formation of choice preference. The positivity of the correlation suggests that participants who perceived the color-change delay as being shorter showed a greater preference index for the reward with the color-change delay. Taken together, our results highlight that the subjective duration of the delay is a crucial factor in forming a choice preference for delayed rewards.

### Subjective duration of the color-change delay

Prior behavioral studies demonstrated that elapsed duration was overestimated in situations where participants were exposed to a visual stimulus involving abundant information content and/or that varied frequently, suggesting that experiencing busy visual environments prolongs subjectively perceived time^4, 5, 8^. In the current study, in contrast, participants underestimated the color-change delay. One important distinction between the visual stimuli used in this and prior studies is that spatiotemporal changes in visual content and information were unexpected in prior studies^4, 8, 46^; on the other hand, the current study presented colored panels that were spatially static, and the temporal change in their colors occurred gradually. Thus, participants were able to predict what the changes would be like from moment to moment. Notably, the increased predictability of events that occur over the course of time shortens subjectively perceived duration^1–3^. Thus, the reverse direction of subjective duration bias in the prior and current studies is attributable to the spatiotemporal complexity that affects the unexpectedness of events in the presented stimuli.

Our imaging analysis of the duration judgment task revealed that reduced activity in the vlPFC was associated with greater perceptual bias of the delay duration. Interestingly, this result is consistent with a recent study demonstrating that the same vlPFC region showed greater activity in individuals with accurate time perception^44^. Therefore, both studies suggest that the vlPFC is associated with accurate time perception, and our study also shows that the vlPFC represents subjectively time-modulated value components while a future reward is anticipated.

### Dynamic value representation during the formation trials

The temporal dynamics of brain activity in the formation trials were modeled based on a probability density distribution that reflected participants’ expectations of the upcoming reward outcome. This model was updated from trial to trial using Bayesian learning, as in prior studies^27, 28^.

One significant extension in the current study is that the probability density distribution was modulated by the subjective duration of the delay estimated from the duration judgment task. Specifically, the subjectively perceived rate of elapsed delay was reflected in temporal characteristics of the probability density of future reward attainment. In particular, during a delay period in which the entire duration was perceived as shorter (i.e., the high frequency color-change delay), participants perceived the delay as elapsing more slowly. This situation was modeled as a wide distribution with a lower peak (Figure 3a *left*).

Based on this novel modeling approach, we found that UFR and AU dynamics were associated with temporally evolving brain signals in the VS and vlPFC, respectively. Crucially, in the duration judgment task, such strong dynamic value signals were absent during the delay period in which a future reward was not anticipated (Supplementary Figure 3). Moreover, the dynamic signals of AU in the vlPFC were specifically observed in the reward choice task (Figure 4). Notably, during the delay period, identical visual stimuli were presented in the reward choice task and duration judgment task, but one critical difference was whether participants were awaiting a future reward. Together, these results demonstrate that anticipation of a future reward produces the value-related temporal dynamics in brain activity, a topic that has not been addressed in previous studies^22, 27–29, 43^.

### Anticipatory utility effect in the prefrontal cortex

Our previous studies showed that the temporal dynamics of the AU were associated with the aPFC, a polar region in the prefrontal cortex^22, 27^ (around Brodmann area 10). Reproducing our previous observations, the current study showed that this polar region showed AU dynamics during the delay period of the reward choice trials (Supplementary Figure 6). On the other hand, in the formation trial, the AU dynamics were observed in the vlPFC (around Brodmann area 45/46/47), posteriorly and laterally to the polar region in the aPFC. The anatomically distinct recruitment of the aPFC and vlPFC suggest that the current formation trials involve distinct value-related signals while participants anticipate a future reward.

It should be noted that participants were exposed to panel color changes during the delay period, which was not the case in the previous studies^22, 27^. Additionally, the temporal dynamics of value models in the current study were modulated based on the subjective duration of the delay period. Finally, the vlPFC region showed reduced activity in individuals with greater time perception bias in the current study. Thus, physical changes in external environments and consequent subjective experiences may together yield the dynamic value representations in the vlPFC for future reward anticipation.

In the prefrontal cortex, posterior lateral regions (around Brodmann area 6/44/45) are associated with attentional orientation^47^, response inhibition^48–51^, set shifting^48, 52–57^, and other executive control functions triggered by the perception of goal-relevant stimuli in external environments. In contrast, more anterior regions (around Brodmann area 10/46) are associated with higher functions such as integrative reasoning^58, 59^ and high-level goal representation^52, 60^. Additionally, polar regions (Brodmann area 10) are involved in intrinsically generated representation such as imagination and/or simulation of events that may occur in the future (episodic future thinking: ^61–64^; see also ^22^ for further discussion). It is thus interesting that the current vlPFC region is located between the posterior lateral and polar regions in the prefrontal cortex, and is implicated in both anticipation of a future reward and perception of changes in the external world. Choice preference in intertemporal decision-making may arise from the integration between the experience of physical events occurring in the external environments and intrinsically represented subjective interpretation of these events.

## Methods

### Participants

Participants (N = 27; age range, 19-31 years; 14 female) were right handed and had no history of psychiatric or neurological disorders. Written informed consent was obtained from all participants. All experimental procedures were approved by the institutional review boards of Keio University and the National Institute for Physiological Sciences. Participants received 8,000 yen for their involvement. They were instructed not to drink any liquid for 4 hours before the experiment. The sample size was determined prior to data collection based on behavioral piloting and previous relevant studies^20–22, 27, 28^.

#### Stimuli

##### Delay period stimuli

Two types of visually presented stimuli were used during the delay period. One type involved eight colored panels (Figure 1a *bottom*) whose colors were gradually and continuously changed throughout the delay (Figure 1b *top*; color-change delay). The screen was segmented by 3 × 3 rectangles, and the colored panels were placed on the peripheral rectangles. Then, in the center rectangle, a picture stimulus indicating a delayed reward (Figure 1a) was presented throughout the delay period. One color change cycle involved transitions from [255, 0, 0] (red, green, blue) to [255, 255, 0]; from [255, 255, 0] to [0, 255, 0]; from [0, 255, 0] to [0, 255, 255]; from [0, 255, 255] to [0, 0, 255]; from [0, 0, 255] to [255, 0, 255]; and from [255, 0, 255] to [255, 0, 0]. Stimulus presentation lasted 48 s.

Low, medium, and high frequencies were used (Figure 1b *top*), and were associated with the presentation of one, two, and four color change cycles, respectively, during the 48-s period. The initial color of each panel was pseudorandomized, and the color of each panel was updated every 100 ms. The rate of color change was identical across the eight panels for each frequency. As control stimuli for the color-change delay, a set of colored panels was used whose colors remained unchanged through the delay was used (color-constant delay; Figure 1b *bottom*). The duration of the color-constant delay ranged from 36-60 s depending on previous choices (Supplementary Figure 1; see also Duration adjustment procedure below).

##### Reward stimuli

The current study used commercially available drinks as reward that could be immediately consumed. Before the experiment, participants were provided with a list of the following beverages: apple, orange, and lychee juices; probiotic drinks; barley tea; and water. Each participant was then asked to choose the drink that would serve as their reward. The reward amount was constant throughout all trials in the delayed reward choice task.

##### Apparatus

The E-Prime programs (Psychology Software Tools, Pittsburgh) controlled the task as well as the delivery of liquid rewards via a syringe pump (SP210iw; World Precision Instruments, Sarasota). Liquids from two 60-ml plastic syringes mounted on the pump were merged into one tube and then delivered to the participant’s mouth through a silicon tube. The flow rate of each syringe was set to 0.75 ml/s, and thus the reward flowed continuously at a rate of 1.5 ml/s. Participants were able to control the liquid flow. Reward delivery continued as long as they pressed a button on a box that they held in their right hand; delivery paused if they released the button, and resumed when they pressed the button again.

#### Behavioral procedures

##### Delayed reward choice task

During fMRI scanning, participants performed a choice task for real liquid rewards delayed by dozens of seconds. At the beginning of each trial, two pictures were presented on the left and right sides of the screen (Figure 1a). One picture indicated a delayed reward with the color-change delay (e.g., red circle), and the other indicated another delayed reward with the color-constant delay (e.g., blue star). Participants were unfamiliar with the delay duration and the relationships between the pictures and delayed rewards.

One trial block was consisted of three types of trials (Figure 1c *top*). Because participants were naive to the relationships between picture items and reward experiences prior to each trial block, the first two trials were conducted so they would form choice preferences concerning delayed rewards by learning these associations (formation trials)^27^. In the first trial, participants were forced to choose the picture indicating the color-change or -constant delay (first formation trial), and choice stimuli were presented until their response. When participants pressed the button corresponding to the picture, the delay period started and colored panels appeared (Figure 1a *bottom*). After the delay period ended, a visual message indicated that the reward, and participants consumed a 6-ml liquid reward. After they consumed the entire amount, the picture choice was replaced by a fixation cross. Then, two identical pictures were presented again, and participants were asked to choose the picture that they selected in the immediately preceding choice. They then received feedback on their choice depending on whether it was correct or incorrect (probe test; Figure 1c *bottom left*). This probe trial was intended to ensure that participants learned the relationships between choice pictures and rewards.

The next trial started after an inter-trial interval. Two identical pictures were presented, and participants were forced to choose the other picture that were not selected in the first formation trial (second formation trial). Immediately after participants pressed the button corresponding to their choice, the colored panels appeared. After the delay period ended, participants consumed a 6-ml liquid reward, then engaged in another probe test for the picture that they chose in the immediately preceding choice. The order of the color-change and -constant delays was kept constant within participants and counter-balanced across participants.

After participants performed the two formation trials, two identical pictures were again presented on the screen. In the third trial, participants chose one of the two options based on their experiences in the two formation trials, as they preferred (choice trial). Depending on their choice, participants received the same delayed reward as in the formation trial. After reward consumption, participants were asked whether they chose the intended choice option (choice confirmation; Figure 1c *bottom left*).

One trial block consisted of two formation trials and one choice trial, and participants performed nine total blocks during fMRI scanning. A message alerted participants before each block began. Each change frequency was used once per scanning run, and one run involved three trial blocks.

Before fMRI scanning, participants received instructions for the task using a computer display. Participants were told that one picture indicated a delayed reward with a color-change delay and the other picture indicated a delayed reward with a color-constant delay. They were asked not to count the duration covertly during the delay period. They were told that the amount of the reward they would receive was identical in all trials. To familiarize participants with the task, one practice trial block was performed (change frequency: 1.5).

We used 18 unique pictures (6 shapes × 3 colors), and each picture was used in only one block. Participants were told that pictures used in one task block were unrelated to those in other task blocks. Picture items used in the practice block were not used in scanning blocks.

The duration of a single trial was 90 s in the formation and choice trials, regardless of participants’ choices in the choice trials^20–22, 27^ (Figure 1c *bottom left*). Thus, the duration of one trial block, the entire duration of one session, and the total amount of liquid received were not dependent on the choices that participants made.

##### Duration judgment task

After the delayed reward choice task, participants performed another task in which they made a judgment about the duration of the color-change and -constant delays (duration judgment task; Figure 1c *bottom right*). Stimuli and behavioral procedures were identical to those in the delayed reward choice task except that 1) they did not receive a liquid reward after the delay period; and 2) instead of choosing a delayed reward, they judged which of the delay periods lasted longer. Participants used a button press to indicate their judgment about the duration. A single trial consisted of 70 s for the experience of a delay, and 15 s for the judgment.

##### Duration adjustment procedures

To estimate subjective duration of the color-change delay and choice preference for the delayed reward with the color-change delay, the duration of the color-constant delay was adjusted using the following procedures.

In the delayed reward choice task, the durations of the color-change and – constant delays were 48 s in the first block of each frequency (Supplementary Figure 1a). The duration of the color-constant delay from the second block was adjusted according to the previous choice, like in our previous studies^20–22, 27^. Specifically, if participants had chosen a reward with a color-change delay in the first reward choice trial, then the duration of the color-constant delay was reduced by 8 s; if participants had chosen a reward with a color-constant delay, then the duration of the color-constant delay was increased by 8 s. The duration of the color-constant delay in the third block was adjusted similarly, but the adjustment of the duration was 4 s. Then, the choice preference for the color-change reward was defined as the preference index. Technically, the preference index was equal to 2 s less or more than the duration of the color-constant delay on the last (third) block, depending on whether the color-changed or -constant reward, respectively, had been chosen in that block. This procedure was applied to each of the three color change frequencies.

A similar adjustment procedure was applied to the duration judgment task to estimate the subjective duration of the color-change delay. Specifically, if participants judged that a color-constant delay was longer in a trial block, duration of the color-constant delay was reduced in the next block, and *vice versa* (Supplementary Figure 1b).

##### Imaging procedures

Functional MRI scanning was conducted on a whole-body 3T MRI system (Siemens Verio, Germany). Functional images were acquired using multi-band accelerated echo-planar imaging [repetition time, 800 ms; echo time, 30 ms, flip angle, 45 deg; slice thickness, 2mm; in-plane resolution, 2 × 2 mm; multi-band factor, 8; 72 slices; matrix size, 96 × 96]. Whole-brain scanning with high temporal resolution allowed us to perform whole-brain exploratory analyses of temporal dynamics during each delay period with sufficient scanning frames. Each run involved 1070 volumes (14.5 min) in the reward choice task, and 630 volumes (8.5 min) in the duration judgment task. For each task, three functional runs were performed (total six runs and 5100 volumes). The 10 initial volumes of each run were excluded from imaging analysis to take into account the equilibrium of longitudinal magnetization. High-resolution anatomical images were acquired using an MP-RAGE T1-weighted sequence [slice thickness: 1mm; in-plane resolution: 1 × 1 mm^2^].

##### Image preprocessing

Functional images were preprocessed using the FSL suite (http://www.fmrib.ox.ac.uk/fsl/; ver. 5.0.8). All functional images were first temporally aligned across the brain volumes, corrected for movement using rigid-body rotation and translation, and then registered to the participants’ anatomical images. The anatomical images were transformed into a standardized MNI template using linear and non-linear registrations implemented in *flirt* and *fnirt*. Functional images were then registered to the standard template by applying registration parameters estimated for the anatomical images. The data were then resampled into 2-mm isotropic voxels, and spatially smoothed with a 5-mm full-width at half-maximum Gaussian kernel.

To minimize motion-derived artifacts due to consumption of liquid rewards, functional images were further preprocessed by GLM estimations with motion parameters and MRI signal time courses (cerebrospinal fluid, white matter and whole brain), and their derivatives and quadratics as nuisance regressors^27, 28, 65, 66^. Then residual of the nuisance GLM was used for standard GLM estimations to extract the task event-related brain activity described below, based on *fsl_regfilt* implemented by the FSL suite.

#### GLM

##### Single level analysis

A GLM approach was used to estimate trial event effects. Parameter estimates were performed by *feat* implemented by the FSL suite. Events of particular interests were the temporal characteristics (dynamics) of brain activity during the delay periods of the formation and choice trials.

The temporal dynamics of the delay-period brain activity were represented by two value representation models, as theoretically forecasted and empirically demonstrated in previous literature^13, 22, 27–29, 38–40^. One model was the AU, reflecting the utility of waiting for a future reward^22, 28, 38–40^, and the other was the UFR, indicating the monotonic increase of the upcoming future reward value as the delay elapsed^13, 22, 28, 38, 41^. The AU model showed a peak value at the beginning of delay and a gradual decrease throughtout the delay, while the UFR model had an inverse temporal pattern^22, 27, 28^.

Events of non-interest consisted of the choices of the formation trials, probe tests, reward consumption in the formation trials, choices in the choice trials, reward consumption in the choice trials, choice confirmation probe, and message presentation regarding the next trial block; these events were coded separately, convolved with the canonical HRF, and included in the GLM.

##### Formation trial

One key issue in modeling delay-period dynamics during the formation trials was that participants were unfamiliar with the upcoming delayed reward because they had no previous experience with it (see also Behavioral procedures)^27, 28^. Nonetheless, it was possible to expect when the future reward became available based on past trials they had experienced in prior blocks. Thus, the current model assumed that prior to the formation trial, participants had an expectation of the delay duration, and this expectation was updated according to experiences gained during the formation trials.

The current analysis modeled the expectation of duration and its updates based on a Bayesian inference learning approach^27, 28, 42^. We assumed that the expectation was expressed by a probability density distribution of reward outcome, and the distribution was calculated as a function of delay time using a gamma function. Then, the distribution was updated every time participants completed a formation trial.

Bayesian inference was performed as

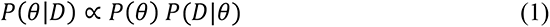

where *D* represents delay duration and *θ* is a parameter that determines probability density. The probability density function (*PDF*) was then calculated as

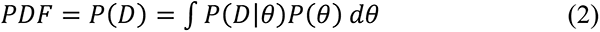

where *P*(*D*|θ) takes a gamma distribution.

If the color-change delay was perceived as shorter or longer than the color-constant delay, the probability density distribution should be modulated in accordance with the subjectively perceived duration. For example, if the color-change delay was be perceived as being shorter, the subjective course of time elapsed more slowly during the color-change delay than during the color-constant delay. Therefore, the probability distribution should be wider along the temporal axis, with lower peak probability. Conversely, if the color-change delay was perceived as longer, the distribution should be temporally narrower with a higher peak. Thus, the probability density function in the color-change delay trial was modulated by the subjective duration of the delay. Specifically, the probability density function was multiplicated by the relative subjective duration, and its time constant (i.e., speed of temporal change) was divided by the relative subjective duration, which was formulated as

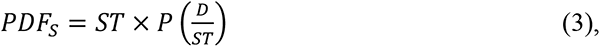

where *ST* is the subjective duration of individual participants for each change frequency as estimated based on the duration judgment task (Figure 3a *left*), and is normalized by the duration of the color-constant delay (i.e., 48 s). Thus, *ST* < 1 if the color-change delay was perceived as being shorter than the color-constant delay, and *ST* ≥ 1 otherwise. In the color-constant delay trial, *ST* was set to 1.

In order to model UFR dynamics during the delay period, the probability density function was integrated from the start of the delay to time *t*,

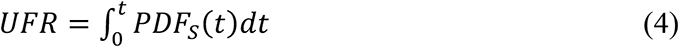

(Figure 3a *middle*). The rationale behind this integration is that, as the delay period elapsed, participants’ expectation of a reward outcome continued to increase. The cumulative probability during the delay reflects the value of the upcoming future reward, although participants did not know exactly when the reward would become available. Then, the AU model was defined as

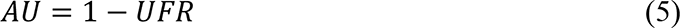

(Figure 3a *right*), as in prior models^22, 27, 28^. The UFR and AU models were calculated for each formation trial of each participant, and then convolved with the canonical HRF (Supplementary Figure 2a).

##### Reward choice trial

For the choice trials, to model the temporal characteristics of value signals during the delay period, the AU and UFR components were defined based on linear functions (Supplementary Figure 2b)^27, 39^. The delay period started after participants chose one of the two alternatives, and there was then a delay with the same duration as the one participants experienced immediately before their choice. The maximal value (1) of the AU model prepared at the beginning of the delay period, and it gradually decreased to the minimum value (0) at the end of the delay. The UFR model had the inverse dynamics. Then, the two dynamic value components were convolved with canonical HRF (Supplementary Figure 2b).

##### Duration judgment task

In the analysis of subjective duration (Figure 5a), we analyzed imaging data obtained during the duration judgment task, rather than the delayed reward choice task, because the former is more appropriate to examine subjectively perceived duration. In GLM, the delay period was modeled using a block effect from the onset to the offset of the delay with a constant value through the delay, but not using the AU or UFR model because this analysis is unrelated to reward value signals during anticipation of a future reward. Other events of non-interest were modeled similarly to the GLM analyses above.

##### Group level analysis

Parameter estimation maps for the AU and UFR models in the formation and choice trials were collected from all participants, and were subjected to a group-level one-sample t-test based on permutation methods (5000 permutations) implemented in *randomise* in the FSL suite. Then voxel clusters were identified using a voxel-wise uncorrected threshold of P < .01, and the voxel clusters were tested for significance with a threshold of P < .05 corrected by the FWE rate. This non-parametric permutation procedure was validated to appropriately control the false-positive rate^67^. The peaks of significant clusters were then identified and listed in tables. If multiple peaks were identified within 12 mm, the most significant peak was retained.

In the analysis of subjective duration (Figure 5a), we restricted exploration within the vlPFC regions showing AU effects during the delayed reward choice task to test whether the AU-related vlPFC regions (Figure 3b *bottom*) are associated with subjective perception of time (see Results). We note that the AU-related vlPFC mask was created based on the imaging analysis of the reward choice task, independently of the data evaluated for the duration judgment task. Then, using this vlPFC mask, statistical corrections were performed using the non-parametric permutation procedure.

##### Data and code availability

The datasets and code supporting the current study are available from the corresponding author (Koji Jimura) on reasonable request.

## Acknowledgments

This work was supported by JSPS Kakenhi (21H05060, 20K07727, 17K01989, and 26350986 to K.J.); NIPS Cooperative Study Program (20-639, 19-635, 18-633, 17-6229, and 21-544 to K.J.); ABiS (16A-073-M02 to K.J.); and AMED (JP21dm0207086 to J.C.).

## Supplementary Figure

**Supplementary Figure 1.**
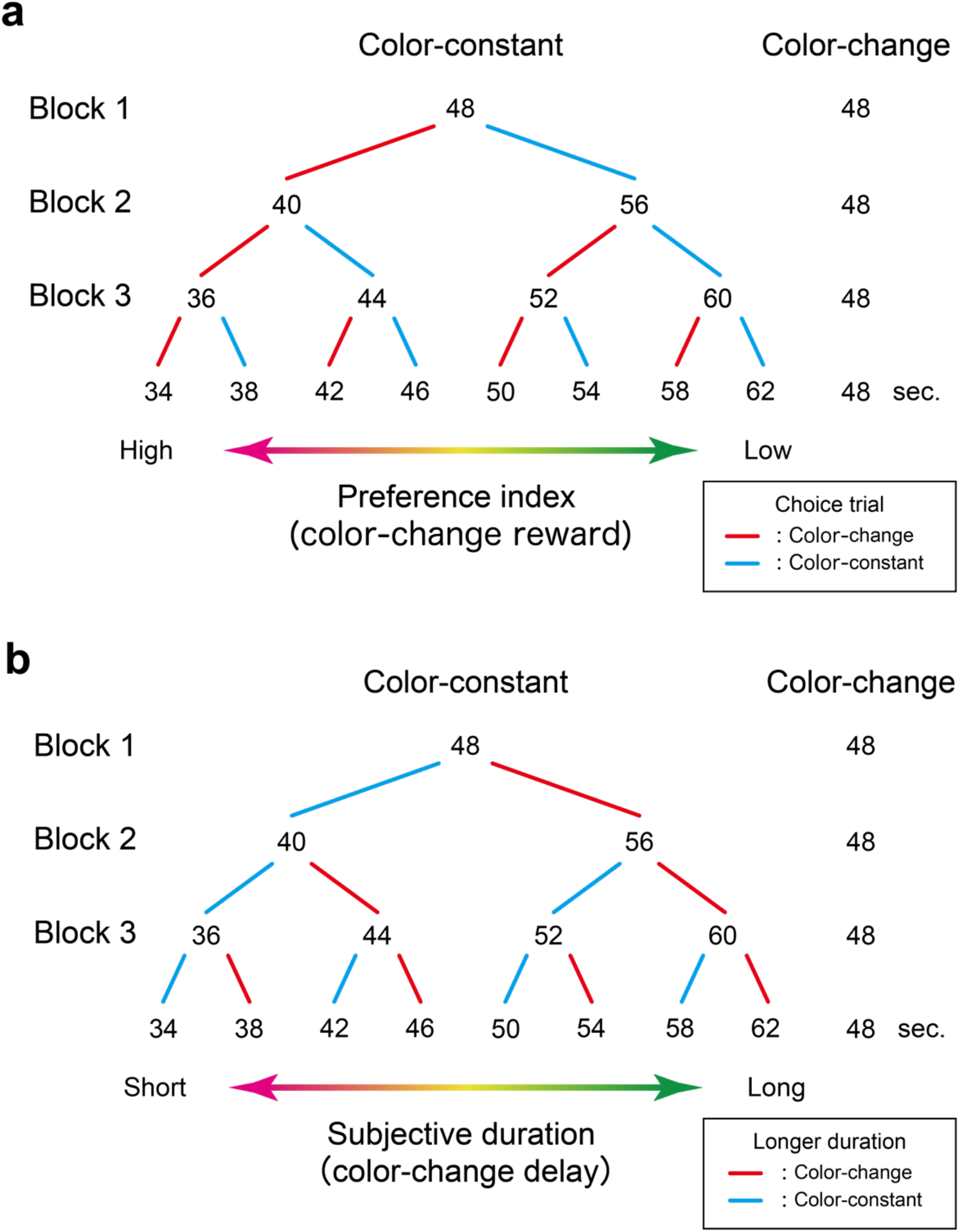
Procedure for adjusting the duration of the color-constant delay. **a**, In the reward choice task, the choice preference index was estimated by increasing or decreasing the duration of the color-constant delay based on the choices in previous choice trials. Then the preference index of the color-change delay reward was estimated as an equivalence of the duration of the color-constant delay. **b**, In the duration judgment task, the subjective duration of the color-change delay was adjusted and estimated similarly based on the judgment which of the delays was longer.

**Supplementary Figure 2.**
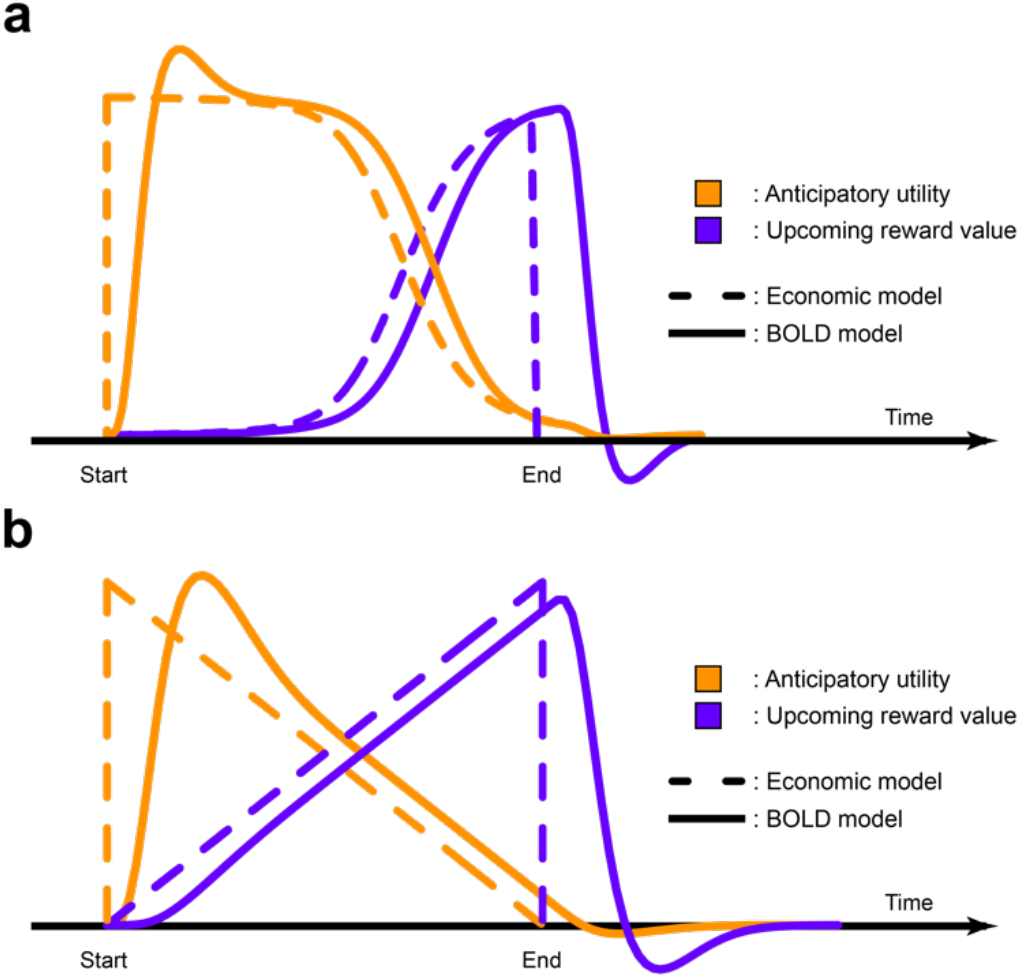
Economic and fMRI models for dynamic value components during delay periods. UFR and AU models were convolved with the canonical HRF to calculate BOLD models in the formation trials (**a**) and the choice trials (**b**). Broken lines indicate economic models estimated based on Bayesian inference learning, and solid lines indicate BOLD models.

**Supplementary Figure 3.**
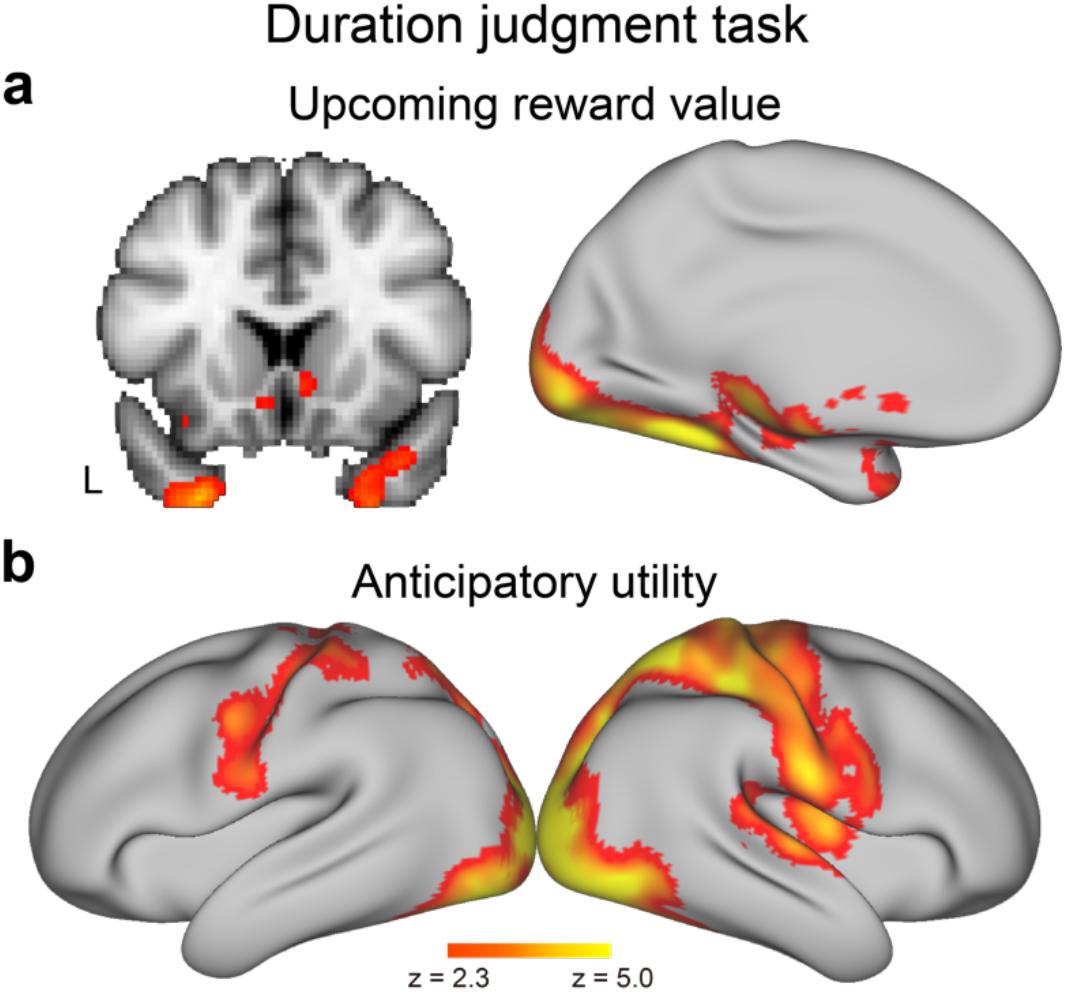
Brain regions showing dynamic value signal components during the delay period of the duration judgment task. Note that in this task, rewards were not delivered and thus participants did not anticipate a reward during the delay period. **a**, UFR effect. **b**, AU effect. Formats are similar to those in Figure 3b.

**Supplementary Figure 4.**
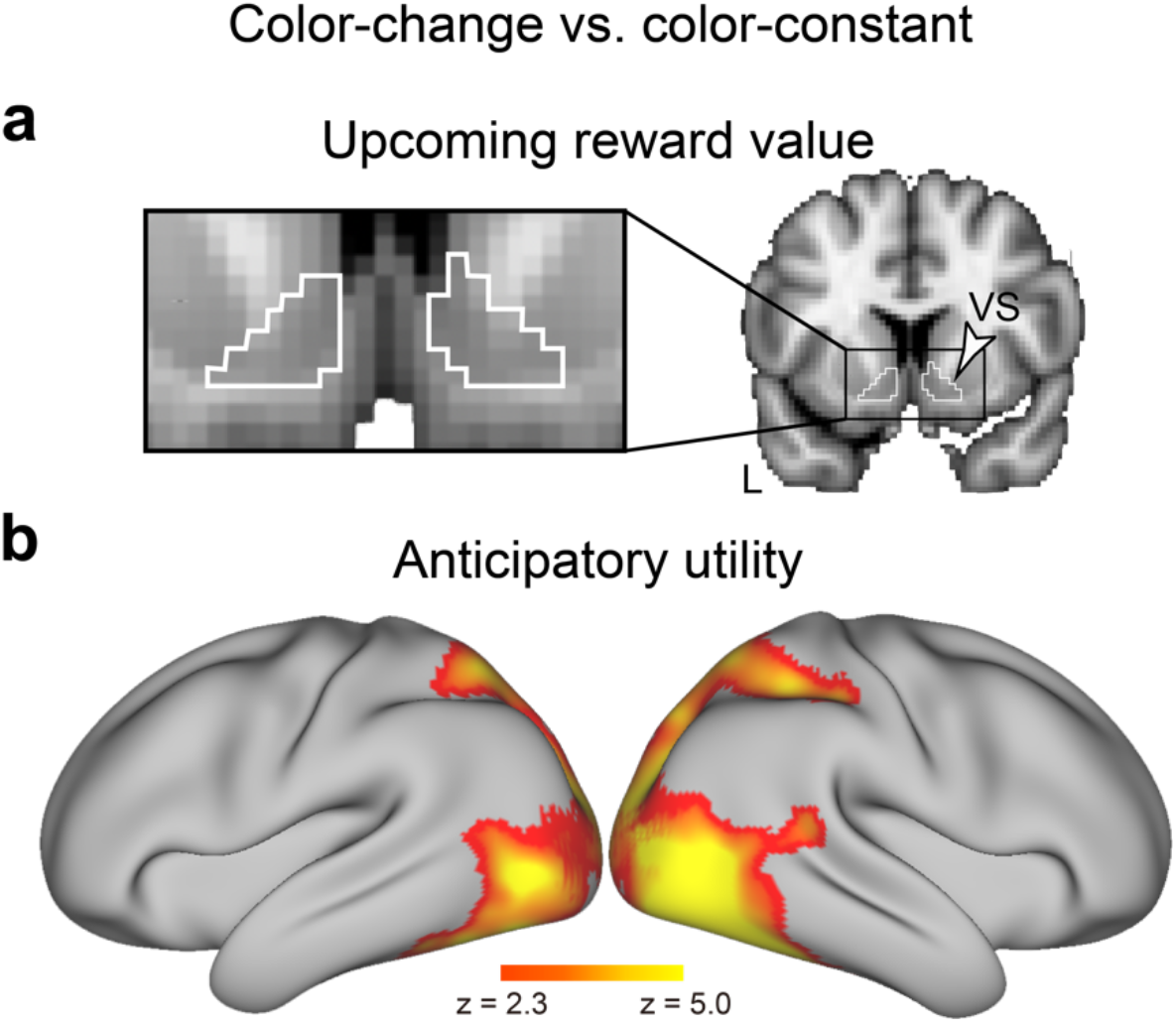
Brain regions showing differential dynamic value components between the color-change and -constant delays in the delayed reward choice task. **a**, UFR effect. **b**, AU effect. Formats are similar to those in Figure 3b.

**Supplementary Figure 5.**
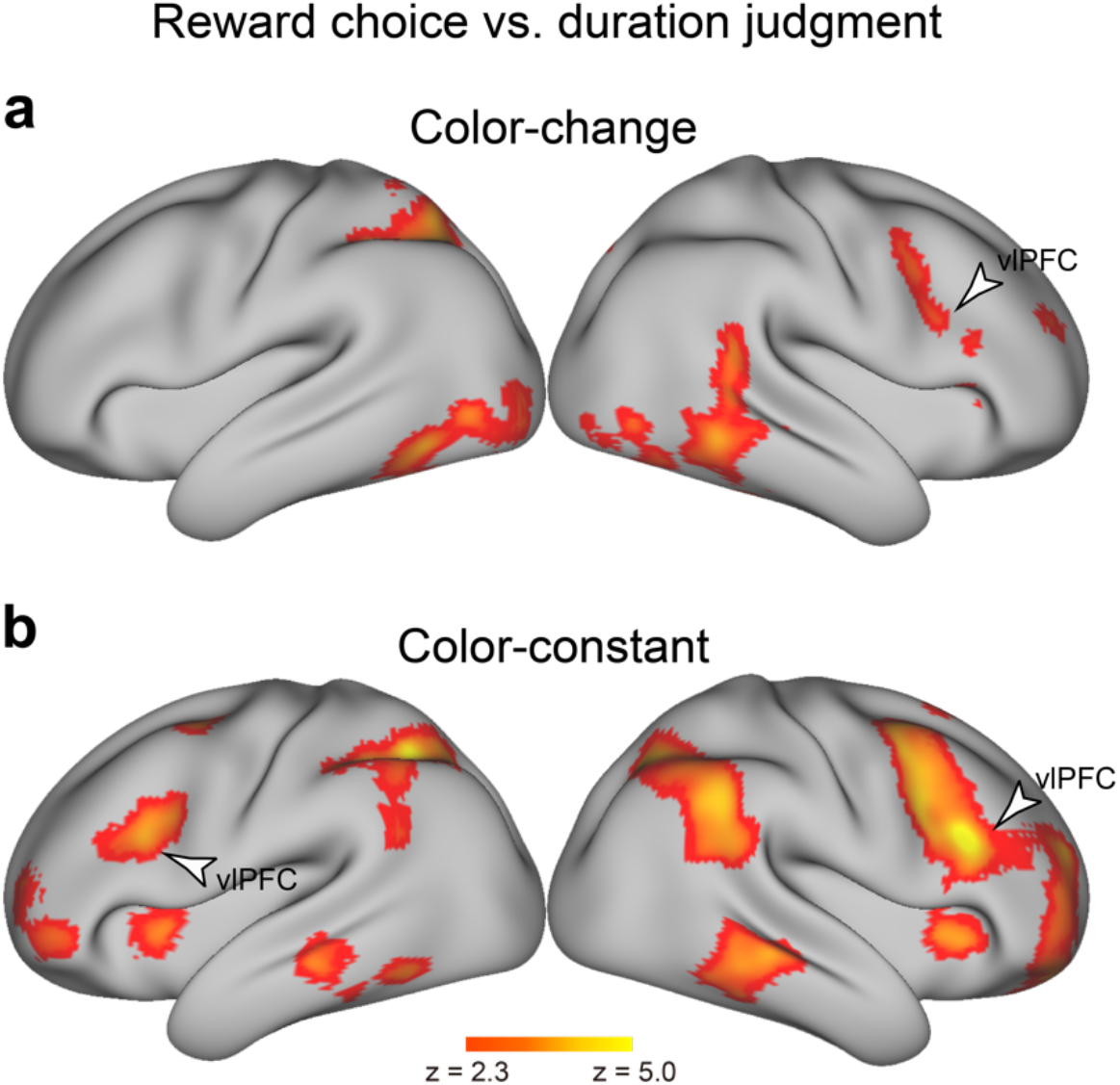
Brain regions showing differential AU effects between the reward choice and duration judgment tasks. **a,** Color-change delay. **b,** Color-constant delay. Formats are similar to those in Figure 3b.

**Supplementary Figure 6.**
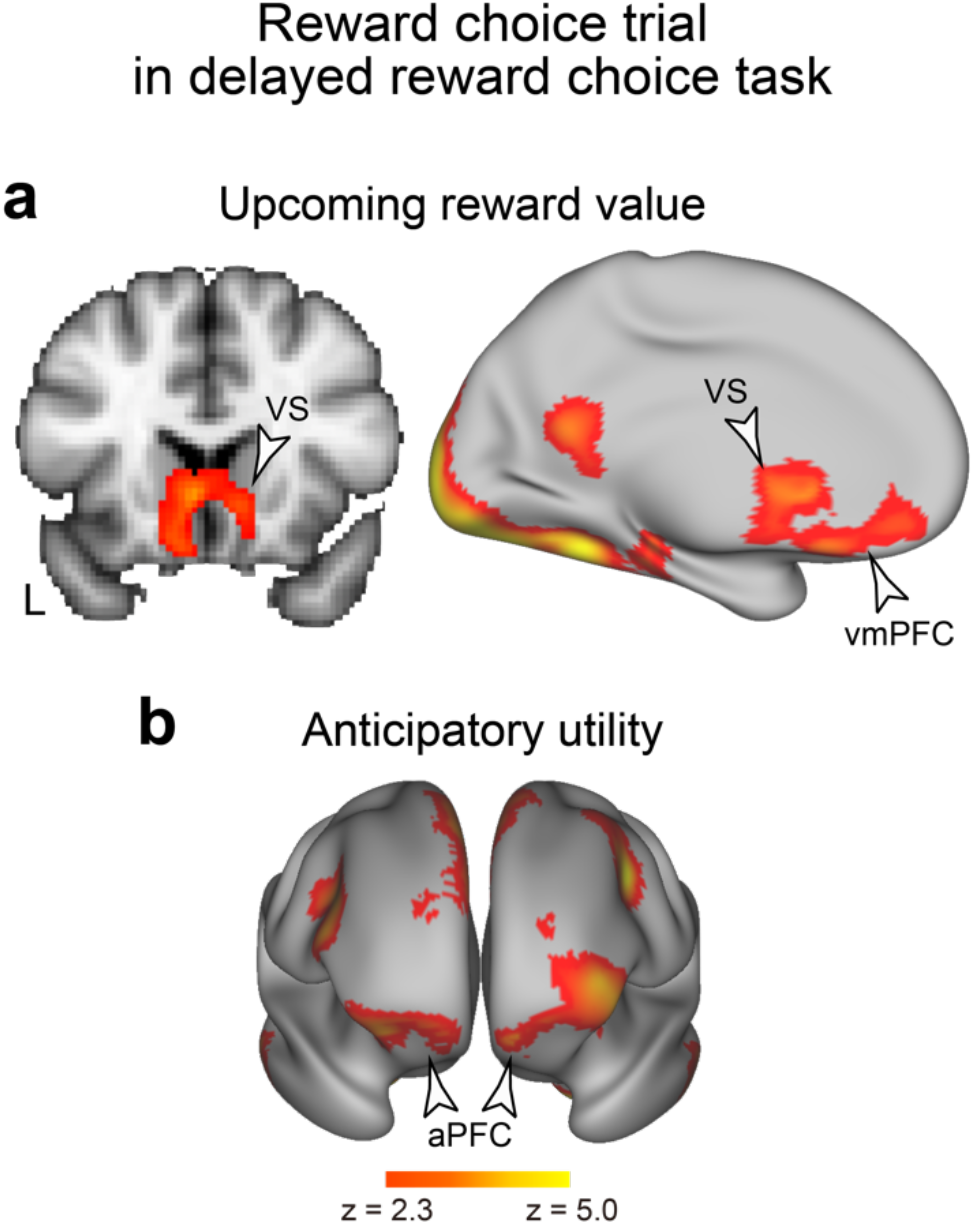
Brain regions showing dynamic value signal components during the delay period of the choice trials in the delayed reward choice task. **a**, UFR effect. **b**, AU effect. Formats are similar to those in Figure 3b.

## Supplementary tables

**Supplementary Table 1.**
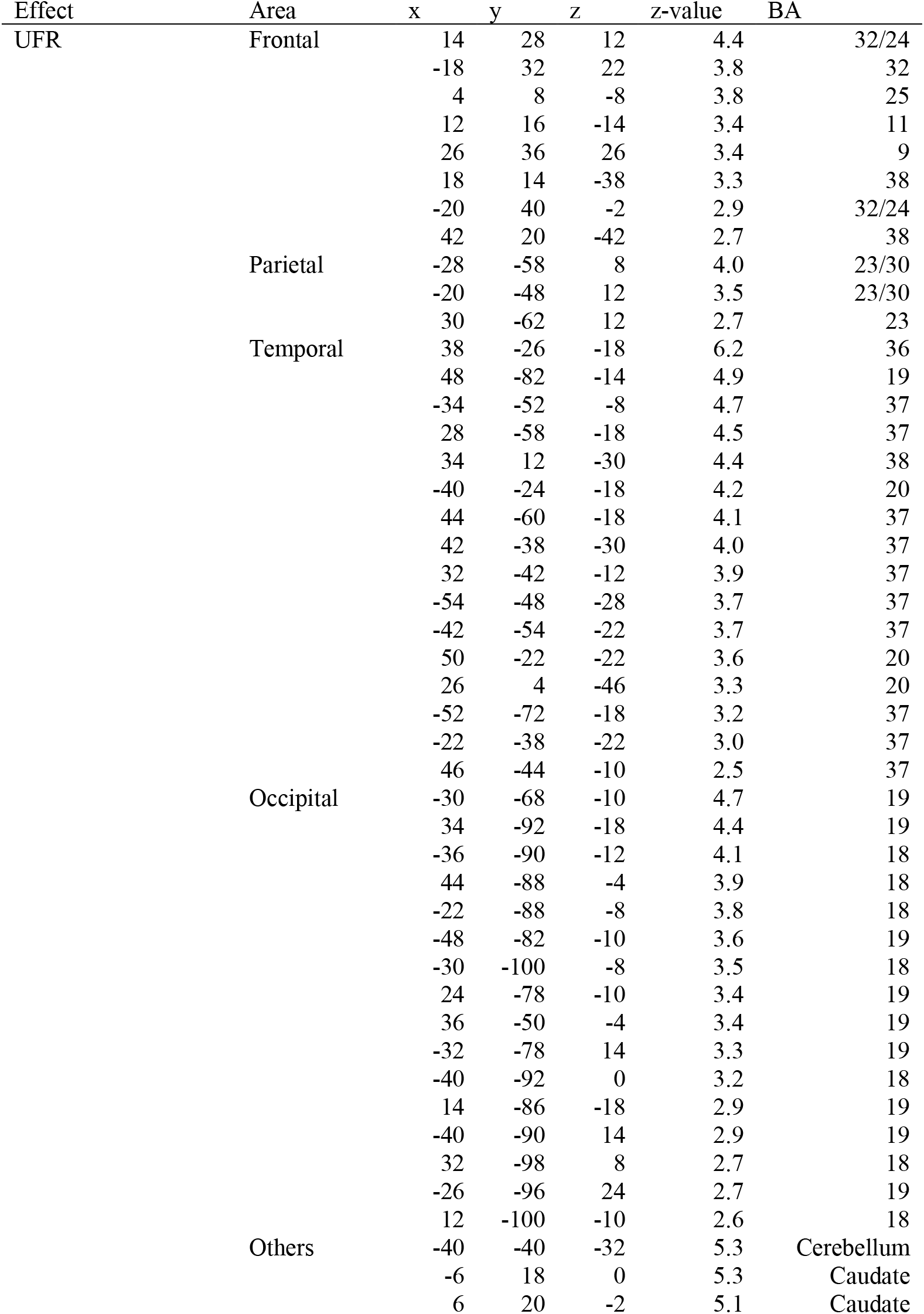
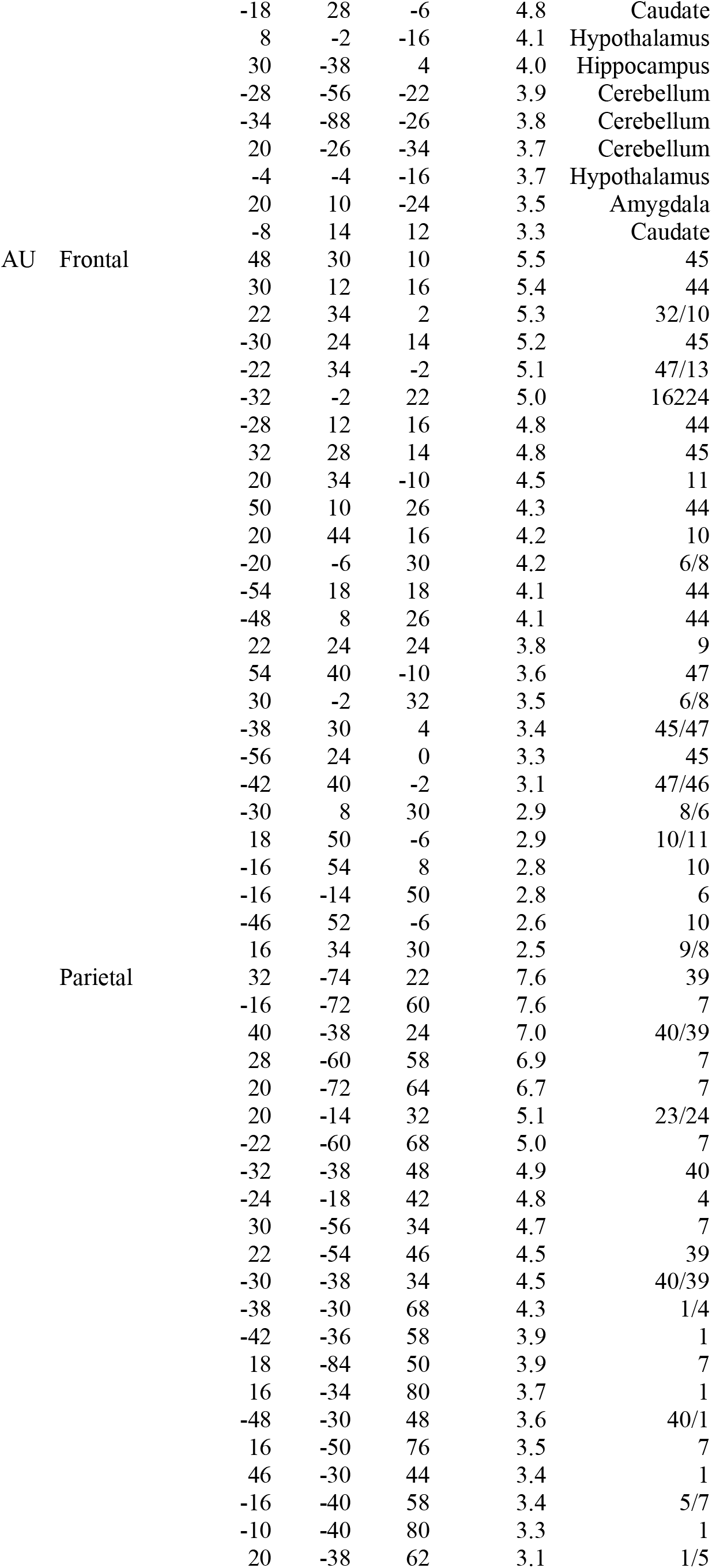
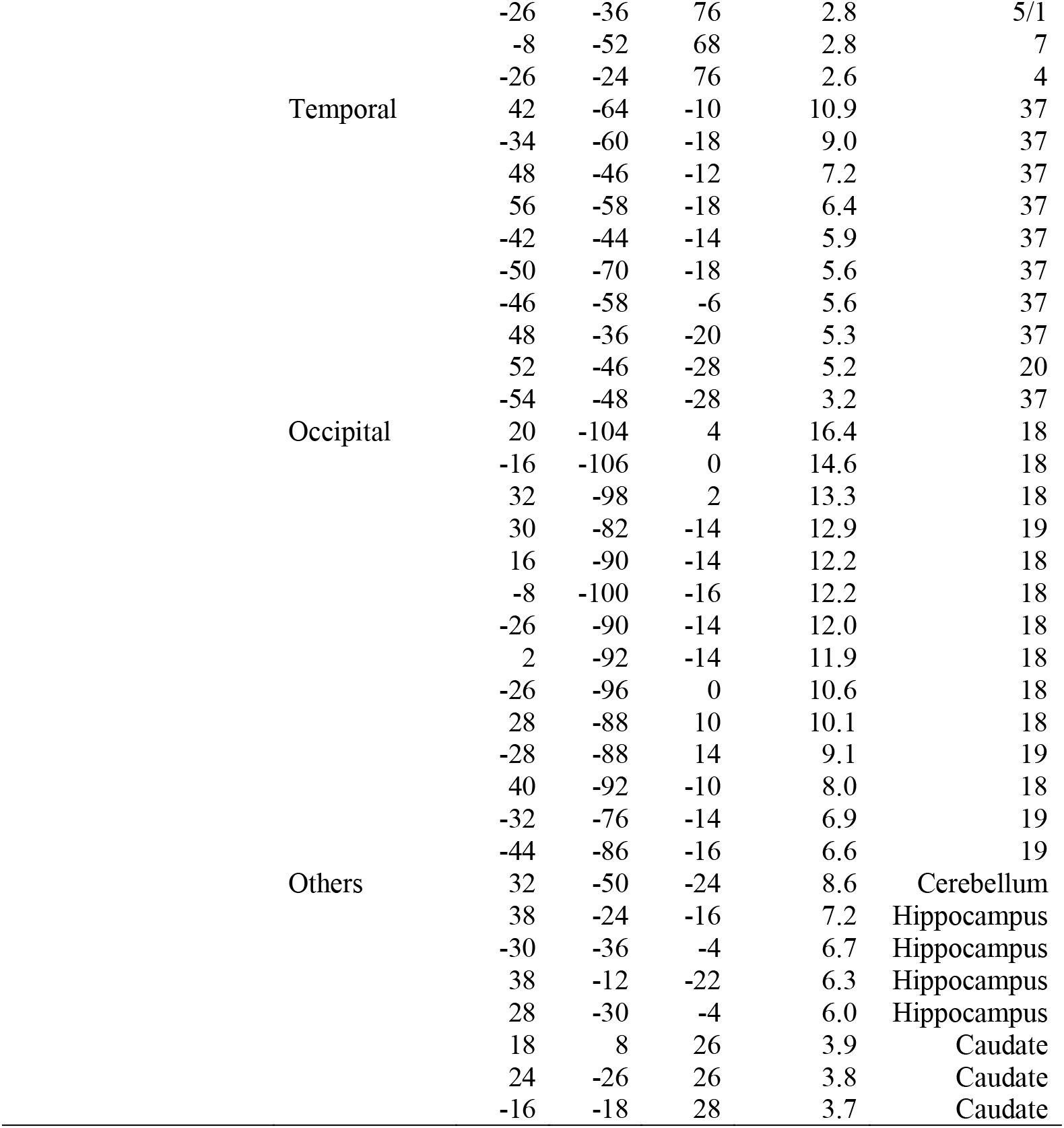
Brain regions showing a significant effect of UFR and AU during the delay period of the formation trial in the reward choice task. Coordinates are listed in MNI space. BA indicates the Brodmann area near the coordinates and is approximate.

**Supplementary Table 2.**
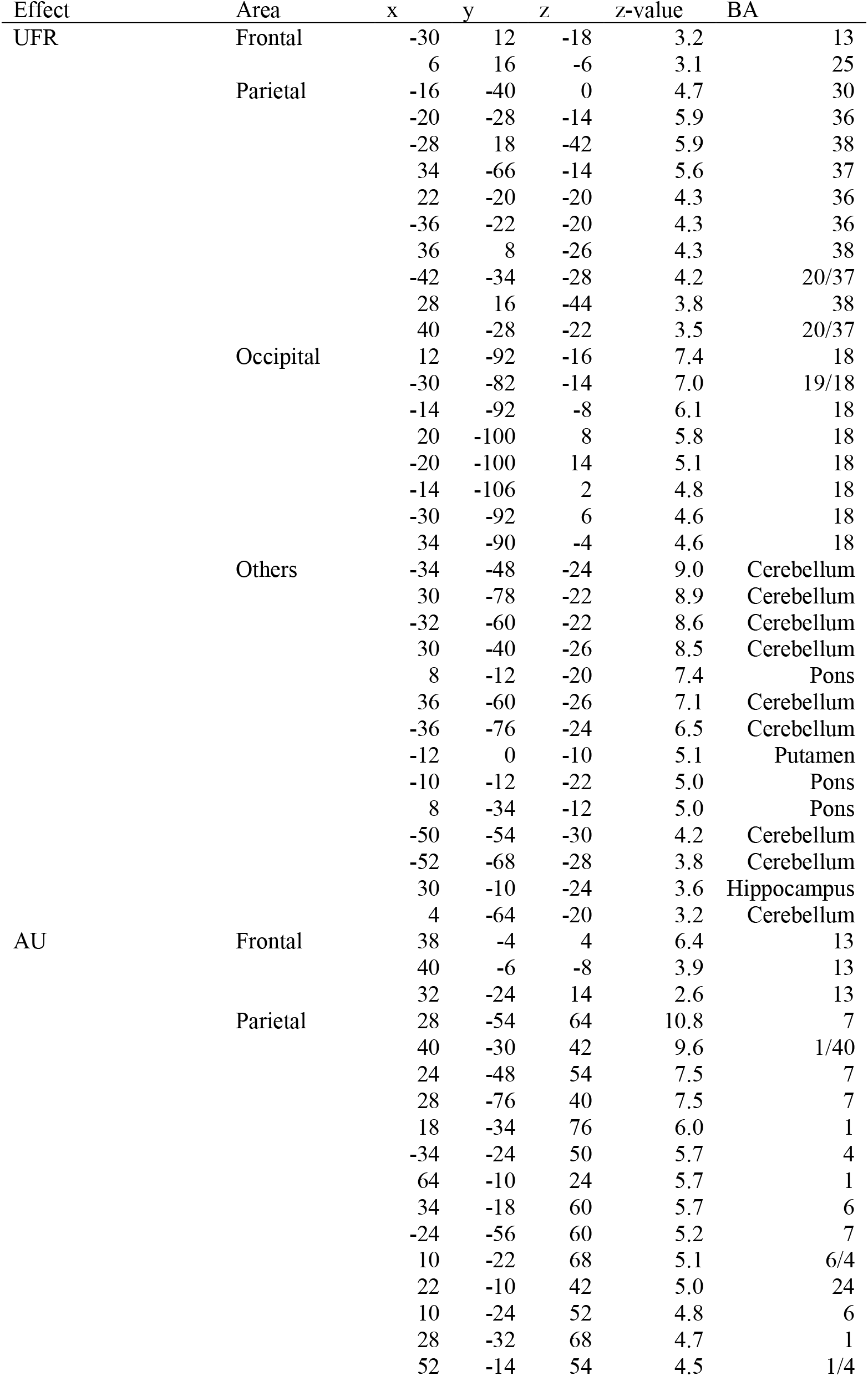
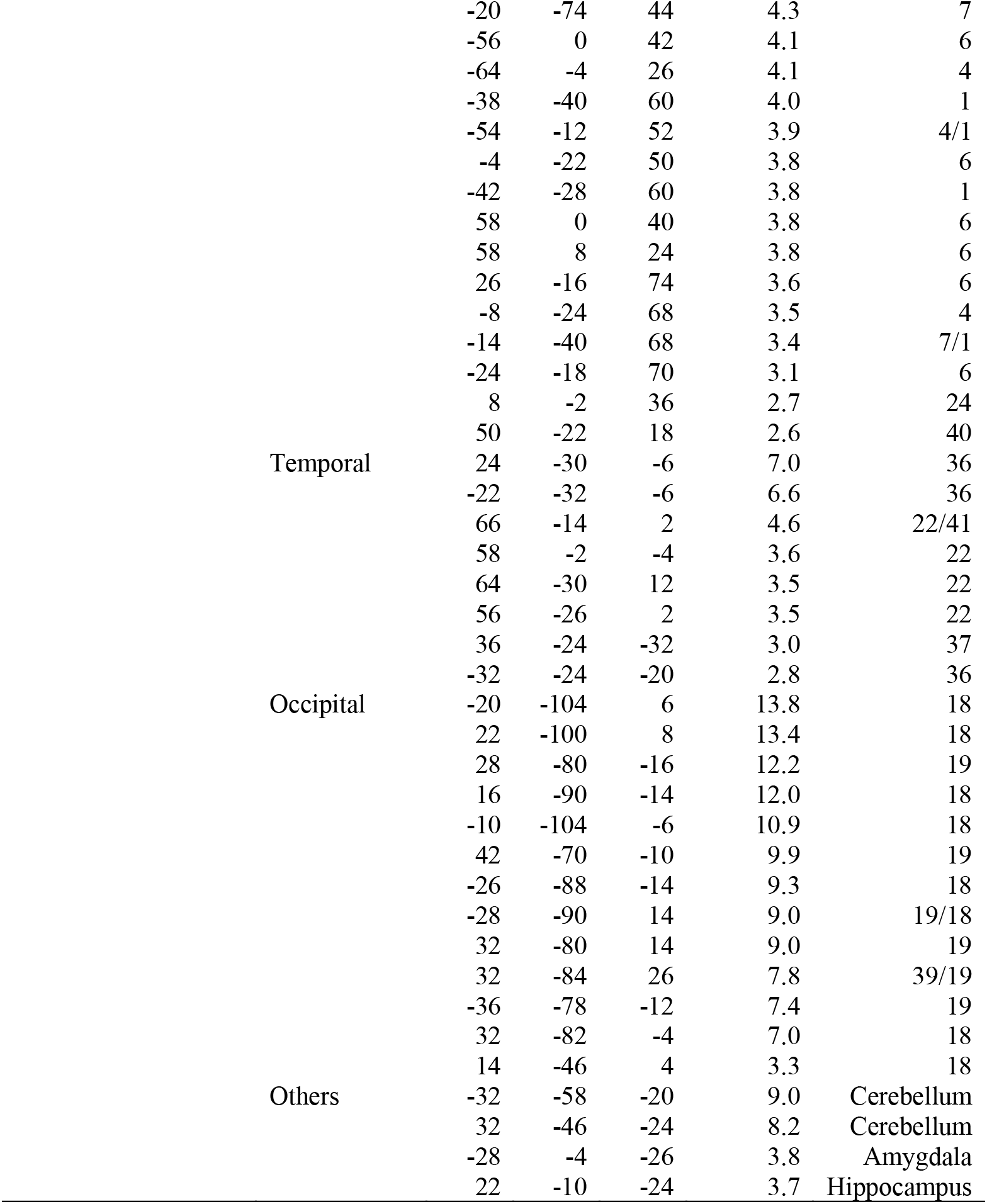
Brain regions showing a significant effect of UFR and AU during the delay period of the formation trial in the duration judgement task. Formats are similar to those in Supplementary Table 1.

| Effect | Area | x | y | z | z-value | BA |
| --- | --- | --- | --- | --- | --- | --- |
| UFR | Frontal | -30 | 12 | -18 | 3.2 | 13 |
|  |  | 6 | 16 | -6 | 3.1 | 25 |
|  | Parietal | -16 | -40 | 0 | 4.7 | 30 |
|  |  | -20 | -28 | -14 | 5.9 | 36 |
|  |  | -28 | 18 | -42 | 5.9 | 38 |
|  |  | 34 | -66 | -14 | 5.6 | 37 |
|  |  | 22 | -20 | -20 | 4.3 | 36 |
|  |  | -36 | -22 | -20 | 4.3 | 36 |
|  |  | 36 | 8 | -26 | 4.3 | 38 |
|  |  | -42 | -34 | -28 | 4.2 | 20/37 |
|  |  | 28 | 16 | -44 | 3.8 | 38 |
|  |  | 40 | -28 | -22 | 3.5 | 20/37 |
|  | Occipital | 12 | -92 | -16 | 7.4 | 18 |
|  |  | -30 | -82 | -14 | 7.0 | 19/18 |
|  |  | -14 | -92 | -8 | 6.1 | 18 |
|  |  | 20 | -100 | 8 | 5.8 | 18 |
|  |  | -20 | -100 | 14 | 5.1 | 18 |
|  |  | -14 | -106 | 2 | 4.8 | 18 |
|  |  | -30 | -92 | 6 | 4.6 | 18 |
|  |  | 34 | -90 | -4 | 4.6 | 18 |
|  | Others | -34 | -48 | -24 | 9.0 | Cerebellum |
|  |  | 30 | -78 | -22 | 8.9 | Cerebellum |
|  |  | -32 | -60 | -22 | 8.6 | Cerebellum |
|  |  | 30 | -40 | -26 | 8.5 | Cerebellum |
|  |  | 8 | -12 | -20 | 7.4 | Pons |
|  |  | 36 | -60 | -26 | 7.1 | Cerebellum |
|  |  | -36 | -76 | -24 | 6.5 | Cerebellum |
|  |  | -12 | 0 | -10 | 5.1 | Putamen |
|  |  | -10 | -12 | -22 | 5.0 | Pons |
|  |  | 8 | -34 | -12 | 5.0 | Pons |
|  |  | -50 | -54 | -30 | 4.2 | Cerebellum |
|  |  | -52 | -68 | -28 | 3.8 | Cerebellum |
|  |  | 30 | -10 | -24 | 3.6 | Hippocampus |
|  |  | 4 | -64 | -20 | 3.2 | Cerebellum |
| AU | Frontal | 38 | -4 | 4 | 6.4 | 13 |
|  |  | 40 | -6 | -8 | 3.9 | 13 |
|  |  | 32 | -24 | 14 | 2.6 | 13 |
|  | Parietal | 28 | -54 | 64 | 10.8 | 7 |
|  |  | 40 | -30 | 42 | 9.6 | 1/40 |
|  |  | 24 | -48 | 54 | 7.5 | 7 |
|  |  | 28 | -76 | 40 | 7.5 | 7 |
|  |  | 18 | -34 | 76 | 6.0 | 1 |
|  |  | -34 | -24 | 50 | 5.7 | 4 |
|  |  | 64 | -10 | 24 | 5.7 | 1 |
|  |  | 34 | -18 | 60 | 5.7 | 6 |
|  |  | -24 | -56 | 60 | 5.2 | 7 |
|  |  | 10 | -22 | 68 | 5.1 | 6/4 |
|  |  | 22 | -10 | 42 | 5.0 | 24 |
|  |  | 10 | -24 | 52 | 4.8 | 6 |
|  |  | 28 | -32 | 68 | 4.7 | 1 |
|  |  | 52 | -14 | 54 | 4.5 | 1/4 |

|  |  |  |  |  |  |
| --- | --- | --- | --- | --- | --- |
|  | -20 | -74 | 44 | 4.3 | 7 |
|  | -56 | 0 | 42 | 4.1 | 6 |
|  | -64 | -4 | 26 | 4.1 | 4 |
|  | -38 | -40 | 60 | 4.0 | 1 |
|  | -54 | -12 | 52 | 3.9 | 4/1 |
|  | -4 | -22 | 50 | 3.8 | 6 |
|  | -42 | -28 | 60 | 3.8 | 1 |
|  | 58 | 0 | 40 | 3.8 | 6 |
|  | 58 | 8 | 24 | 3.8 | 6 |
|  | 26 | -16 | 74 | 3.6 | 6 |
|  | -8 | -24 | 68 | 3.5 | 4 |
|  | -14 | -40 | 68 | 3.4 | 7/1 |
|  | -24 | -18 | 70 | 3.1 | 6 |
|  | 8 | -2 | 36 | 2.7 | 24 |
|  | 50 | -22 | 18 | 2.6 | 40 |
| Temporal | 24 | -30 | -6 | 7.0 | 36 |
|  | -22 | -32 | -6 | 6.6 | 36 |
|  | 66 | -14 | 2 | 4.6 | 22/41 |
|  | 58 | -2 | -4 | 3.6 | 22 |
|  | 64 | -30 | 12 | 3.5 | 22 |
|  | 56 | -26 | 2 | 3.5 | 22 |
|  | 36 | -24 | -32 | 3.0 | 37 |
|  | -32 | -24 | -20 | 2.8 | 36 |
| Occipital | -20 | -104 | 6 | 13.8 | 18 |
|  | 22 | -100 | 8 | 13.4 | 18 |
|  | 28 | -80 | -16 | 12.2 | 19 |
|  | 16 | -90 | -14 | 12.0 | 18 |
|  | -10 | -104 | -6 | 10.9 | 18 |
|  | 42 | -70 | -10 | 9.9 | 19 |
|  | -26 | -88 | -14 | 9.3 | 18 |
|  | -28 | -90 | 14 | 9.0 | 19/18 |
|  | 32 | -80 | 14 | 9.0 | 19 |
|  | 32 | -84 | 26 | 7.8 | 39/19 |
|  | -36 | -78 | -12 | 7.4 | 19 |
|  | 32 | -82 | -4 | 7.0 | 18 |
|  | 14 | -46 | 4 | 3.3 | 18 |
| Others | -32 | -58 | -20 | 9.0 | Cerebellum |
|  | 32 | -46 | -24 | 8.2 | Cerebellum |
|  | -28 | -4 | -26 | 3.8 | Amygdala |
|  | 22 | -10 | -24 | 3.7 | Hippocampus |

**Supplementary Table 3.**
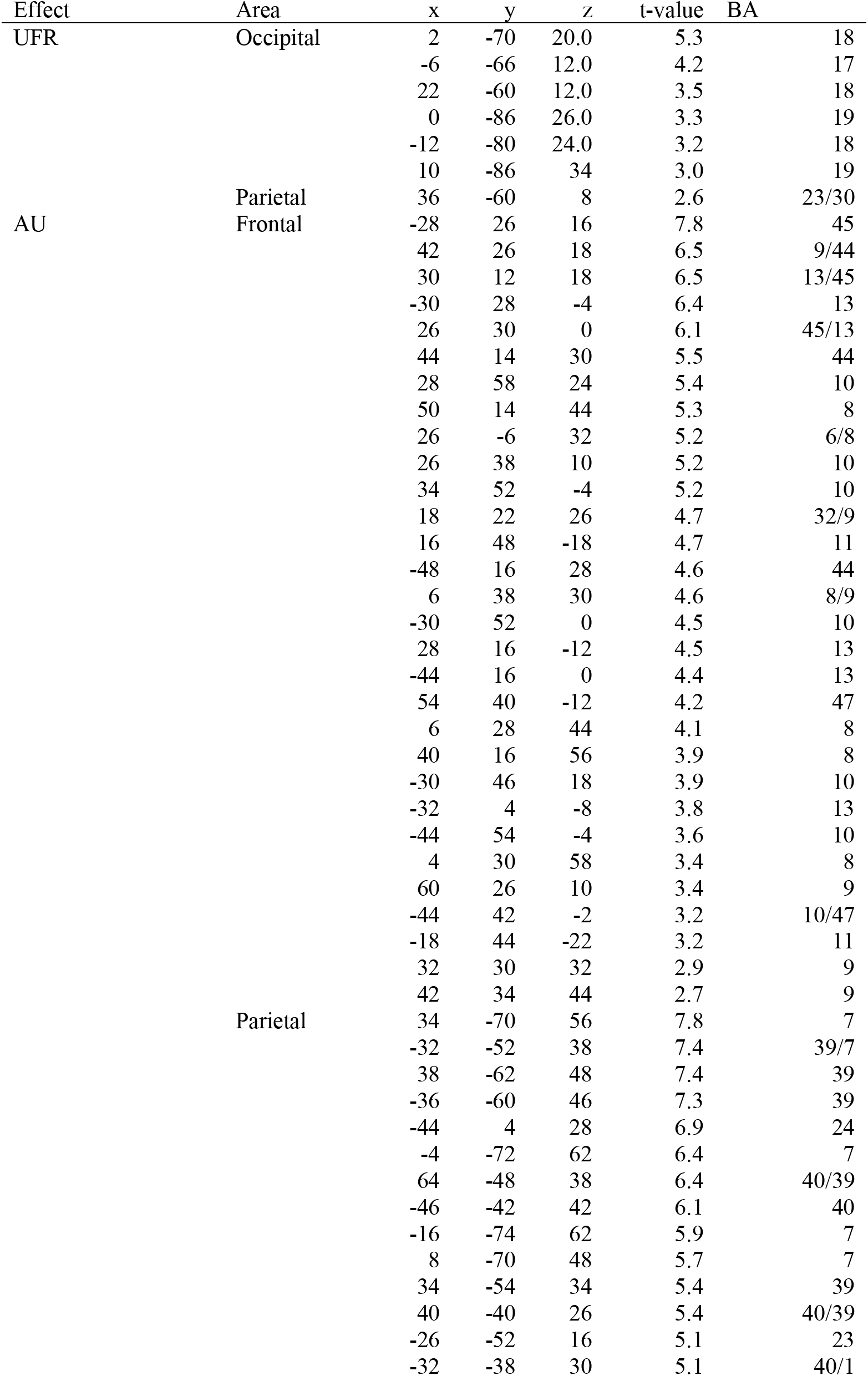
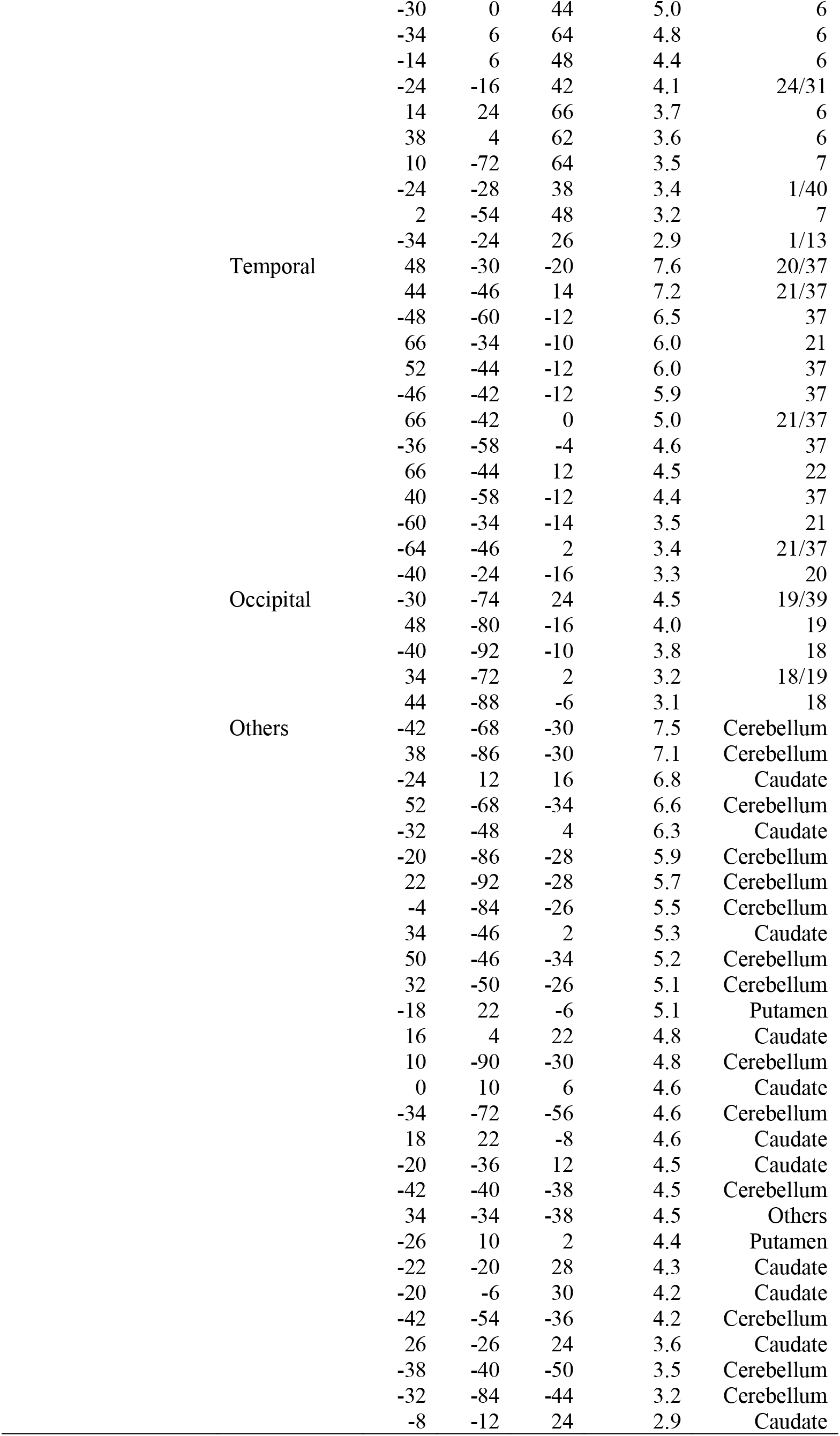
Brain regions showing a significant differential effect of UFR and AU during the delay period of the formation trial between the reward judgment task and the duration judgment task. Formats are similar to those in Supplementary Table 1.

**Supplementary Table 4.**
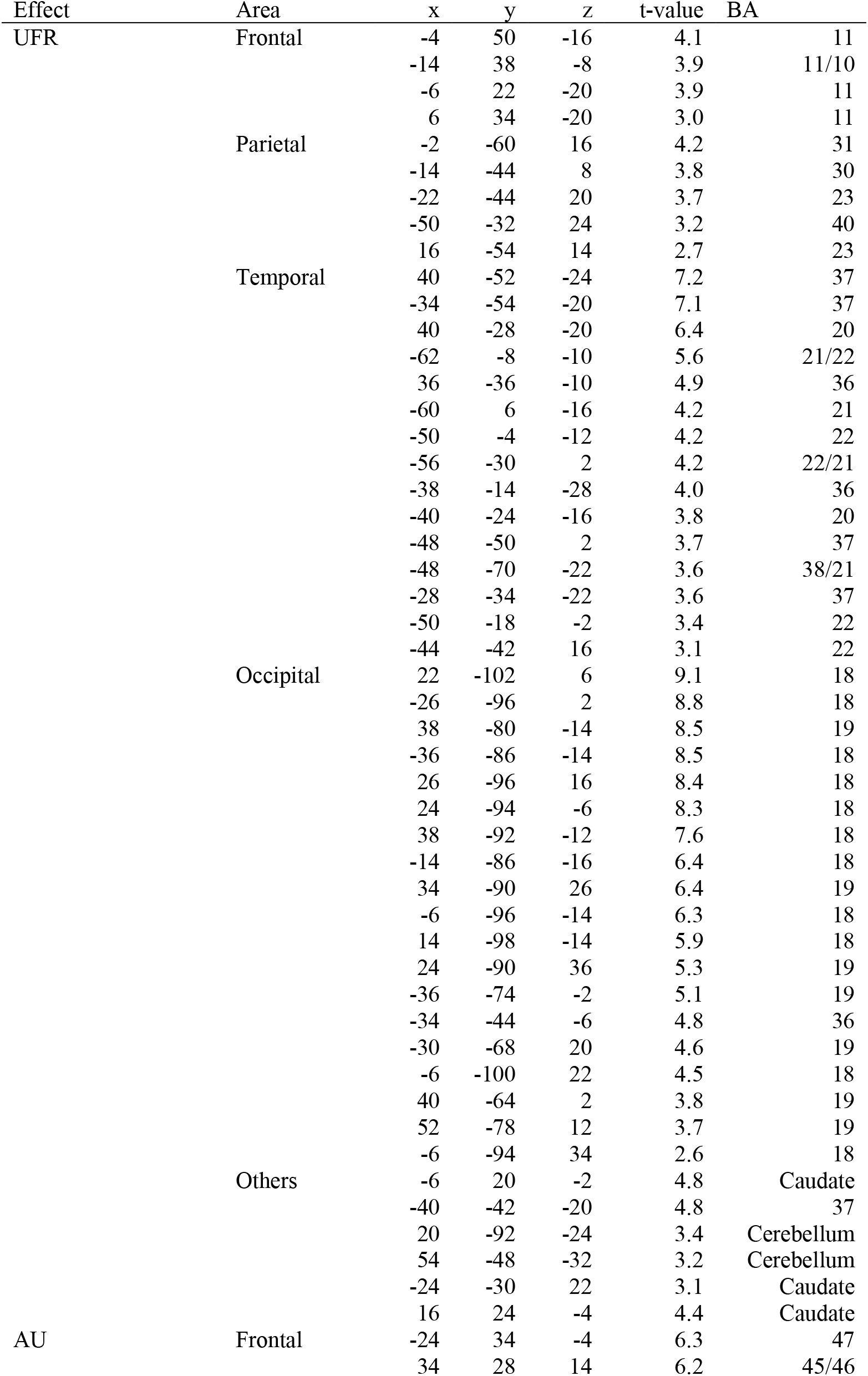
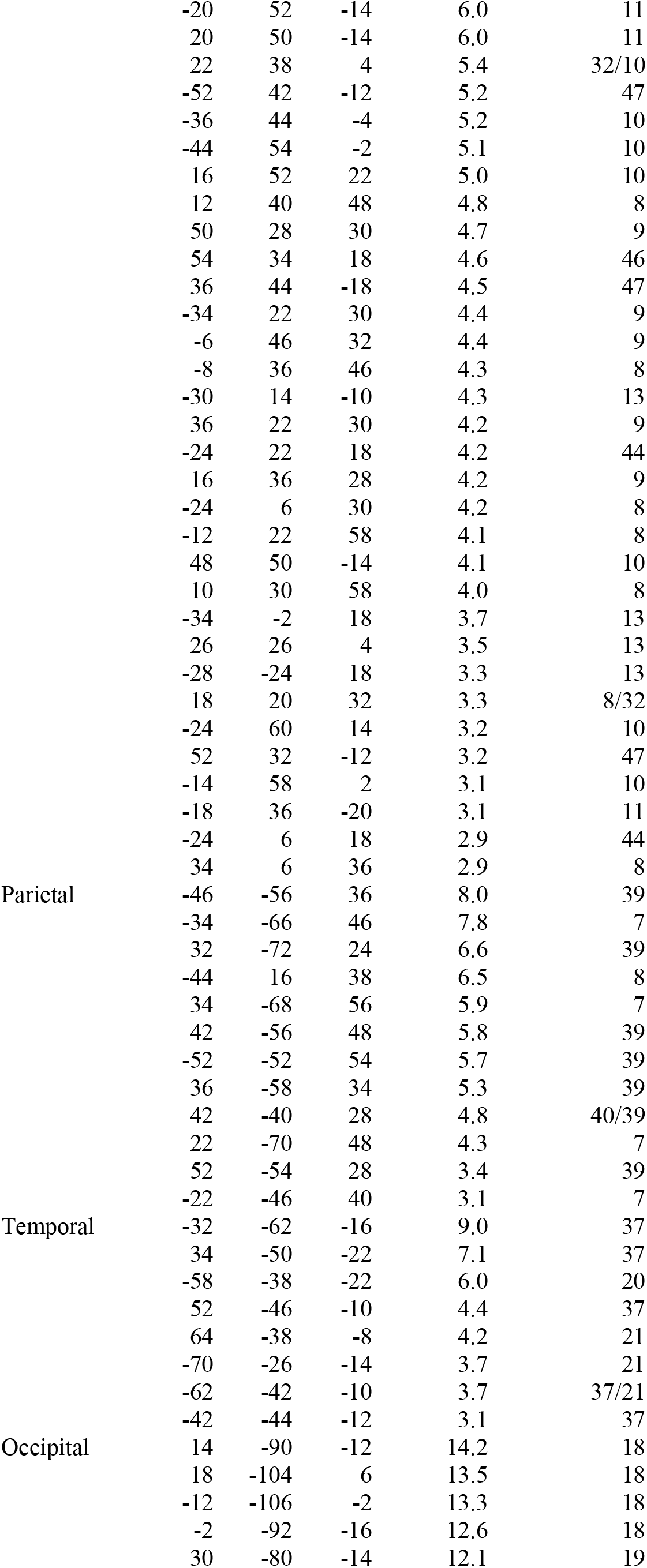
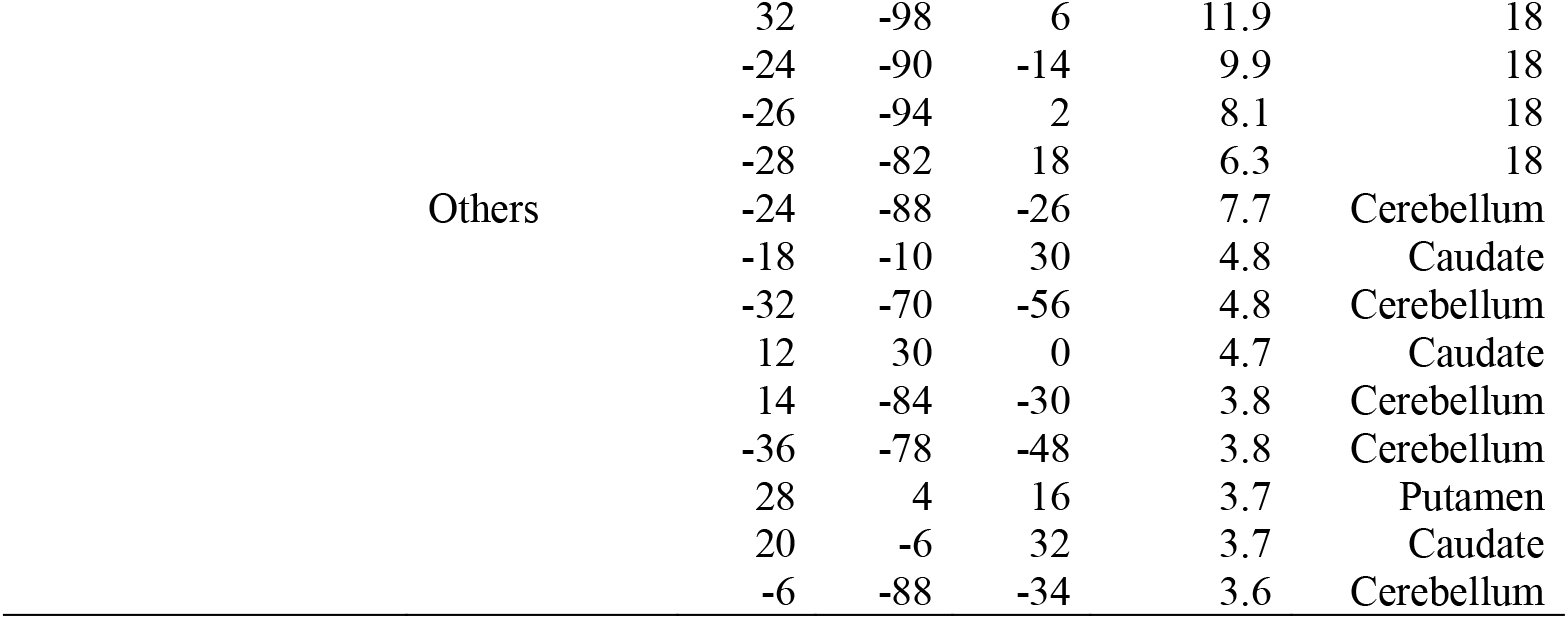
Brain regions showing a significant effect of UFR and AU during the delay period of the choice trial in the reward choice task. Formats are similar to those in Supplementary Table 1.

